# Dopamine regulates the exploration-exploitation trade-off in rats

**DOI:** 10.1101/482802

**Authors:** François Cinotti, Virginie Fresno, Nassim Aklil, Etienne Coutureau, Benoît Girard, Alain R. Marchand, Mehdi Khamassi

**Affiliations:** Institut des Systèmes Intelligents et de Robotique, Sorbonne Universités, UPMC Univ Paris 06, CNRS F-75005, Paris, France; CNRS, Institut de Neurosciences Cognitives et Intégratives d’Aquitaine (INCIA, UMR 5287), Bordeaux, France; Université de Bordeaux, INCIA, Bordeaux, France

**Keywords:** Dopamine, Exploration-exploitation dilemma, Decision-making, Reinforcement learning, Non-stationary bandit task

## Abstract

In a volatile environment where rewards are uncertain, successful performance requires a delicate balance between exploitation of the best option and exploration of alternative choices. It has theoretically been proposed that dopamine controls this exploration-exploitation trade-off, specifically that the higher the level of tonic dopamine, the more exploitation is favored. We demonstrate here that there is a formal relationship between the rescaling of dopamine positive reward prediction errors and the exploration-exploitation trade-off in simple non-stationary multi-armed bandit tasks. We further show in rats performing such a task that systemically antagonizing dopamine receptors greatly increases the number of random choices without affecting learning capacities. Simulations and comparison of a set of different computational models (an extended Q-learning model, a directed exploration model, and a meta-learning model) fitted on each individual confirm that, independently of the model, decreasing dopaminergic activity does not affect learning rate but is equivalent to an increase in exploration rate. This study shows that dopamine could adapt the exploration-exploitation trade-off in decision making when facing changing environmental contingencies.

## Introduction

All organisms need to make choices for their survival while being confronted to uncertainty in their environment. Animals and humans tend to exploit actions likely to provide desirable outcomes, but they must also take into account the possibility that environmental contingencies and the outcome of their actions may vary with time. Behavioral flexibility is thus needed in volatile environments in order to detect and learn new contingencies[1]. This requires a delicate balance between exploitation of known resources and exploration of alternative options that may have become advantageous. How this exploration/exploitation dilemma may be resolved and regulated is still a subject of active research in the fields of Neuroscience and Machine Learning[2]–[5].

Dopamine holds a fundamental place in contemporary studies of learning and decision-making. The role of phasic dopamine signals in supporting learning now appears strongly established[6]–[8]. Dopamine reward prediction error signals have been identified in a variety of instrumental and Pavlovian conditioning tasks[9]–[12]. They affect plasticity and action value learning in cortico-basal networks[13]–[15] and have been directly related to behavioral adaptation in a number of decision-making tasks in humans, non-human primates [16] and rodents[17]–[20]. Accordingly, it is commonly assumed that manipulations of dopamine activity affect the rate of learning, but this could represent a misconception.

Besides learning, the role of dopamine in the control of behavioral performance is still unclear. Dopamine is known to modulate incentive choice (the tendency to differentially weigh costs and benefits) [21], [22] and risk-taking behavior[23], as well as other motivational aspects such as effort and response vigour[24]. Because dopamine is one of the key factors that may encode success or uncertainty, it might modulate decisions by biasing them toward options that present the largest uncertainty[25], [26]. This would correspond to a “directed” exploration strategy[5], [27], [28]. Alternatively, success and failure could affect tonic dopamine levels and control random exploration of all options, as recently proposed by Humphries et al. (2012)[29]. This form of undirected exploration, which is often difficult to distinguish from performance, may be viewed as “noise” in the choice process[30]–[32]. Previous computational analyses of behavioral data in stochastic tasks have yielded mixed results, some suggesting a promotion of random[33] or directed[26] exploration by dopamine with possibly an effect on learning[26], others a reduction of random exploration[29], [34], [35], and yet others an effect on learning only[36].

In the present study, we first show formally that under fairly general assumptions, any manipulation that reduces the magnitude of dopamine positive reward prediction errors does not change learning rate but instead changes the level of random exploration. We then proceed to test this idea experimentally in rats while applying a variety of computational models to the behavioral data. To dissociate learning and performance components, probabilistic tasks where the best option changes with time (known in Machine Learning as non-stationary bandit tasks) are particularly appropriate, because they require periodical exploration and relearning phases and are amenable to computational modeling using well characterized reinforcement learning methods[37]. In this work, we develop such a 3-armed bandit task in rats with varying levels of uncertainty to investigate how dopamine controls the exploration level within an individual. We then examine the effects of dopamine blockade on learning and performance variables following injection of various doses of the D1/D2 receptor antagonist flupenthixol in different sessions. We follow by replicating these data with an extended reinforcement learning model (Q-learning) with forgetting and verify our conclusions on a variety of alternative models such as a directed exploration model, an ε-greedy random exploration model, and a meta-learning model. This allows us to explicitly distinguish learning from exploration variables, and to show that dopamine activity is specifically involved in controlling the level of random exploration rather than the learning rate.

## Results

### Mathematical relationship between reward prediction errors and random exploration-exploitation

We first give here a formal demonstration (**Supporting material**) that in a Q-model such as the one we applied to our task, a reduction of the amplitude of positive reward prediction errors (RPEs) directly translates into an increase in random exploration levels. In other words, it is mathematically equivalent to changing the value of the inverse temperature, and has no effect on the learning rate parameter. Briefly, in this demonstration, we assume that positive, but not negative prediction errors are pharmacologically reduced, which corresponds to a decrease in the value of the reward when it is present. On each trial, the Q-value of the performed action is revised in proportion to the RPE, so it is exactly a fraction of what it would be in the absence of pharmacological manipulation. For non-performed actions, if there is forgetting, the Q-value decreases in proportion to the Q-value itself, which preserves proportionality. As a result, throughout the learning process, all Q-values are downscaled in the same proportion as the reward. When these values are plugged into the softmax process, the result is exactly equivalent to a decrease of the inverse temperature, again in the same proportion. The learning rate or the forgetting rate are not affected in any way. This result shows that under fairly general conditions the effects of a pharmacological manipulation of dopamine-dependent learning should be described as changes in exploration rate rather than as changes in learning rate. Indeed, manipulation of these two factors predicts distinct behavioral profiles: under different learning rates, both performance and win-shift curves (see Methods) should differ at early stages of blocks, but then converge to similar levels (**Supplementary Fig. 2a,b**). Conversely, when only the exploration rate differs, performance curves should tend toward different asymptotes while win-shift curves should shift downwards as a whole when the exploration rate increases (**Supplementary Fig. 2c,d**).

### Dopamine blockade affects exploratory behavior

We then undertook to confirm this result experimentally on a 3-armed bandit task in rats. As a first step, we examined rats’ behavior at different phases of the task once it was well acquired. We focused our analysis on the learning phase that is required each time the target lever changes. Overall, the rats were able to identify the correct lever over the course of a block, despite the stochasticity of rewards (risk).

Performance (**Fig. 1b**) increased within a block toward an asymptote in both low and high risk conditions (F(5,110) = 186.7, p<0.0001), with better performance being observed under low risk (F(1,22) = 148.2, p<0.0001). Dopaminergic blockade by flupenthixol, a D1-D2 dopamine receptor antagonist, decreased performance (F(3,66) = 5.61, p=0.0017) irrespective of trial phase or risk condition (largest F=1.83, p = 0.15). We then looked at exploration, indexed as a first approximation by a win-shift index which only includes shifts from the current best lever (**Fig. 2a**). This index decreased within blocks (F(5,85) = 27.7, p<0.0001) but increased with risk (F(1,17) = 25.7, p<0.0001). Dopaminergic blockade dose-dependently elevated win-shift at all stages (F(3,51) = 14.5, p<0.0001), without interacting with trial or risk (largest F: F(3,51) = 1.40, p = 0.25). Because win-shift in the early stages of blocks may not reflect exploration, but rather a return to a previously reinforced lever, we also limited our analysis of win-shift to the last 8 trials when performance has stabilized, meaning that the correct lever has been identified. In this case also there was a significant dose effect (**Supplementary Fig. 1**). Another index of shifting behavior, lose-shift (**Fig. 3a**), which may denote a correction strategy, was not significantly affected by the pharmacological condition, possibly because of a ceiling effect. We then examined whether the effects of flupenthixol on both win-shift and performance resulted from a negative effect on learning rate, or from a positive effect on exploration rate[38]. In the present task, these two factors predict distinct behavioral profiles, in particular when considering asymptotic behavior (**Supplementary Fig. 2**). Behavioral data indeed point to a change in exploration rate, as individual performance levels during the last six trials of blocks were significantly affected by flupenthixol dose (F(3,66) = 8.85, p < 0.0001) (**Supplementary Fig. 3a,b**). Similarly, the win-shift rate in the last 6 trials of blocks was significantly affected by flupenthixol dose (F(3,66) = 8.76, p < 0.0001) (**Supplementary Fig. 3c,d**). This persistent difference in the curves at the end of each learning phase is strongly suggestive of a change in exploration rate.

**Figure 1.**
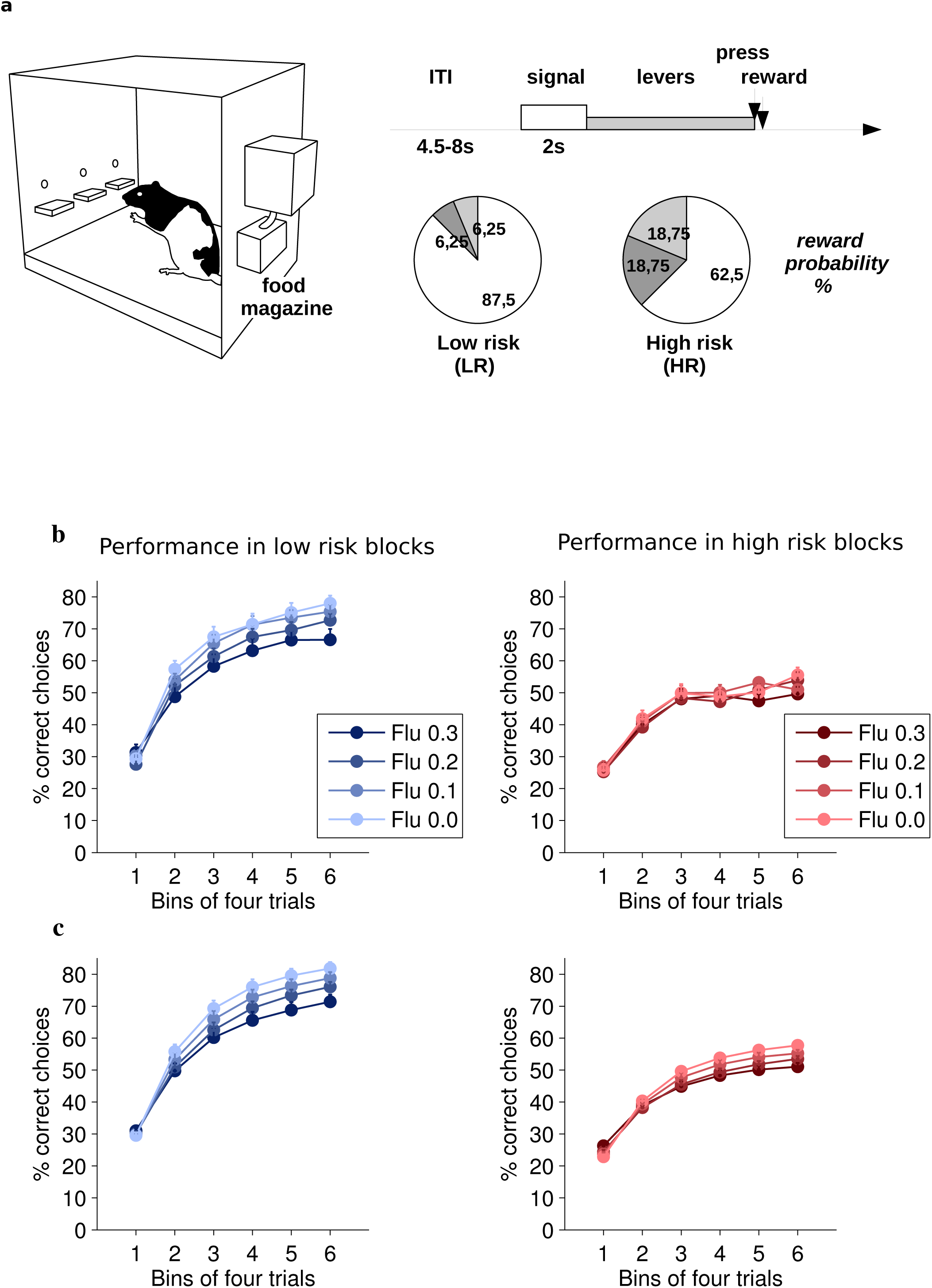
Actual and simulated performances. **(a)** Outline of the experimental task. **(b)** Average formance of rats (n=23) across a block as a function of risk and flupenthixol dose (mean + SEM). Average performance of the model (unconstrained simulations).

**Figure 2.**
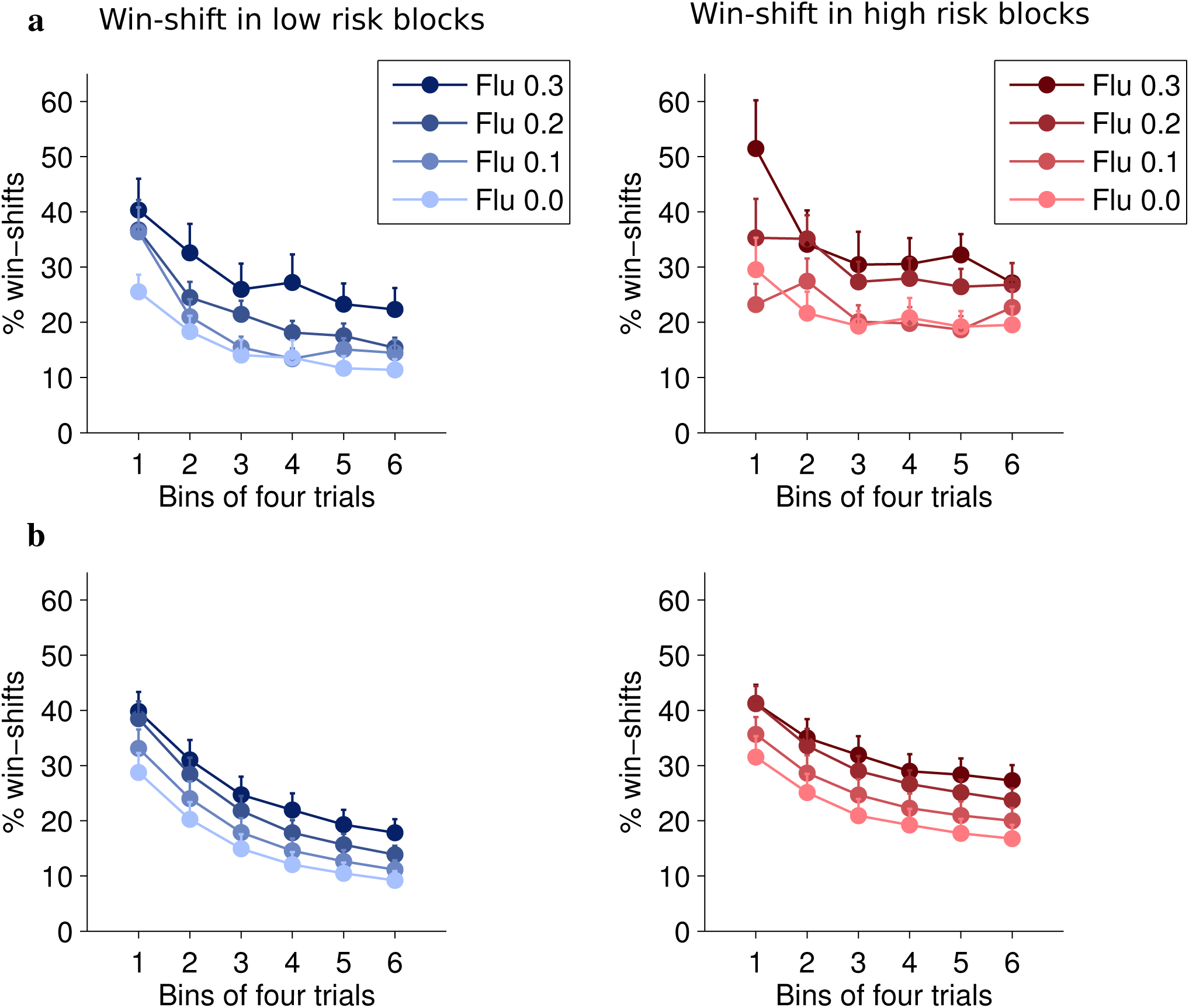
Actual and simulated exploratory choices. **(a)** Average win-shift of rats (n=23) across a block unction of risk and flupenthixol dose (mean + SEM). **(b)** Average win-shift of the model nstrained simulations).

**Figure 3.**
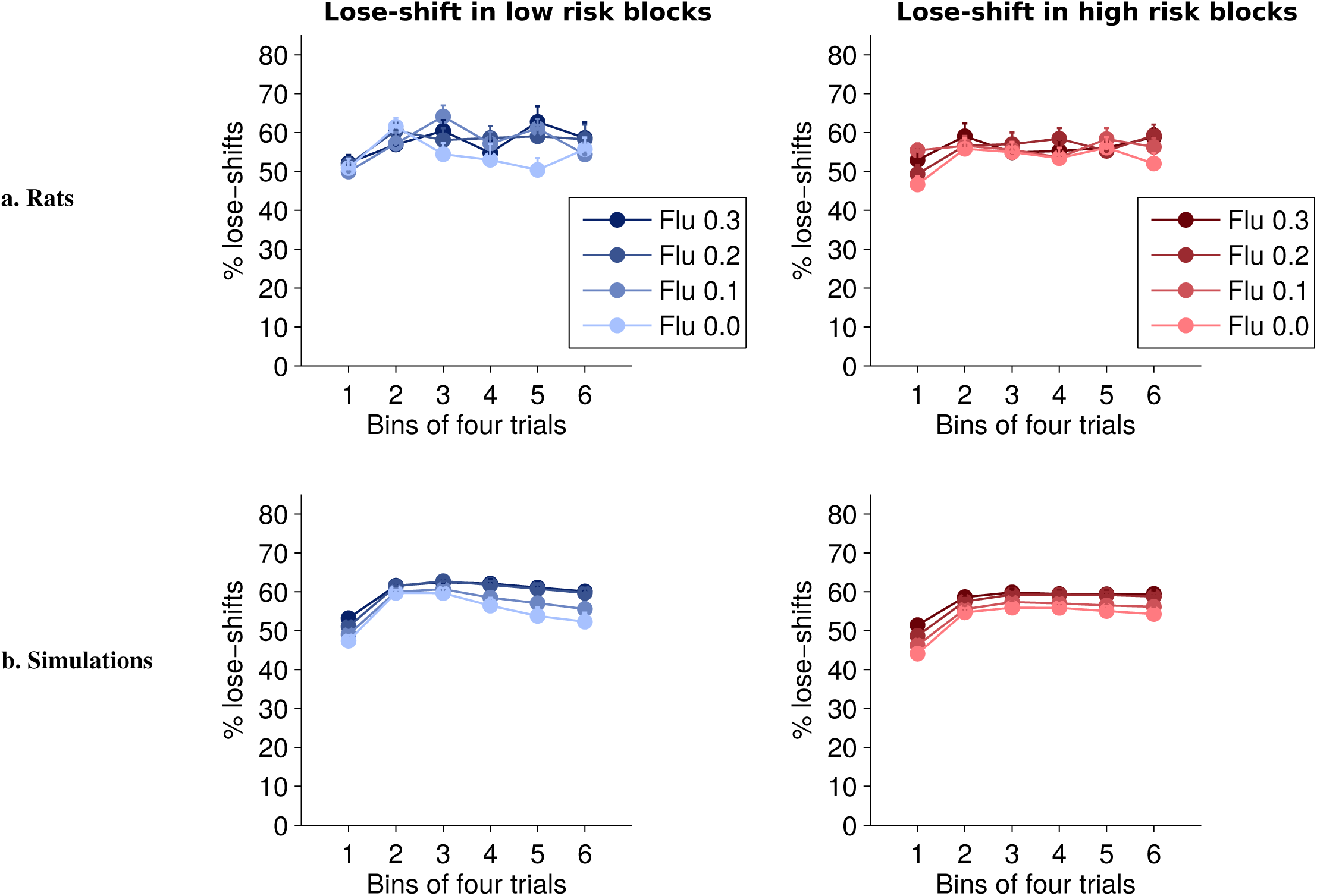
Lose-shift index, i.e. the proportion of trials a rat makes a different choice after an unrewarded trial. **(a)** Average lose-shift of rats (n=23) across a block for different risk and dose conditions (mean +SEM). This index is not significantly affected by risk, and pharmacological conditions (p > 0.11) but a nt trial effect is observed (F(5,110) = 6.83, p < 0.0001). A post hoc test corrected for multiple comparisons using the Bonferroni procedure indicates that lose-shift in the first bin is significantly smaller all other stages of blocks (p<0.0158) except bin number 4 (p=0.17). This effect might be able to persistance, as rats probably continue to press the previous target lever unsuccessfully djusting their behaviour. **(b)** Simulations of the extended Q-learning model reveal a more complex t dynamic with a significant decrease in lose-shift also occuring at the end of blocks. In the simulations, significant main effects of dose (p = 0.0007) and risk (p<0.0001) are also detected. Similarly to win-shift (see **supp. fig. 1** and **7**), simulations of the standard Q-learning model, which are not shown here,stematically overestimate lose-shift levels.

### Flupenthixol alters exploration parameter in simulated rats

We then examined how well a Q-learning model extended with a forgetting mechanism was able to account for behavioral data (see **Methods**). Average action values derived from the model, individually fitted to each rat for each dose of flupenthixol and constrained to the rat’s actual choices, predicted a very high proportion of the variance of individual rats’ choices, (67.7% to 96.4%, **Supplementary Table 1** and **Supplementary Fig. 4**). Moreover, unconstrained simulated data generated using these optimized parameters were highly similar to the actual behavior of the rats (**Fig. 1c, Fig. 2b, Fig. 3B, Supplementary Fig.1, and Supplementary Fig. 6**). When experimental and simulated data were pooled together for repeated-measures ANOVA, there was no significant main effect of simulations on within-block evolution of either performance (Fig. 1c, F(1,22) = 4.27, p = 0.051) or win-shift (Fig. 2b, F(1,17) = 0.006, p = 0.94). However, a significant interaction between simulations and risk did emerge in the case of performance (F(1,22) = 4.42, p = 0.047) and there was also a significant interaction between trials and simulations for both indicators (F(5,110) = 6.98, p<0.0001 for performance; F(5,85) = 2.90, p = 0.018 for win-shift). Crucially however, no interaction involving simulations and flupenthixol dose could be detected (smallest p=0.10). Moreover, when analyzed separately from experimental data, simulated data did in fact replicate the effects of flupenthixol on both measures (F(3,66) = 5.23, p = 0.0027 and F(3,66) = 5.69, p = 0.0016 for performance and win-shift respectively).

### Model optimization dissociates exploration from learning

To disentangle the effects of flupenthixol on learning versus performance, we then examined the values of the different parameters *α*, β and *α*_2_ across pharmacological conditions (**Fig. 4**). Flupenthixol had no discernible effect on the learning rate α (Friedman ANOVA χ^2^(3) = 4.04, p = 0.26) or on the forgetting rate *α*_2_ (χ^2^(3) = 1.38, p = 0.71), but clearly decreased the exploration parameter β (χ^2^(3) = 15.1, p = 0.0018). *Post-hoc* tests revealed that β for 0.2 and 0.3 mg/kg was significantly smaller than for 0 mg/kg (p=0.012 and p = 0.0024 respectively). Thus, the only parameter of the model significantly affected by dopaminergic blockade was the exploration parameter which decreased (i.e. exploration increased) as dopaminergic inhibition increased. We next generalized this result by testing the optimized parameters of a range of different models which include a standard Q-learning model (**Supplementary Fig. 7**), an ε-greedy model of action selection (**Supplementary Fig. 8**), a model of directed exploration (**Supplementary Fig. 9**) which includes an uncertainty bonus to bias decision-making towards options associated with a large value uncertainty[25] and a meta-learning model which sets the value of the inverse temperature based on accumulated reward prediction errors (**Supplementary Fig. 10**). In all of these cases, the only parameter that significantly varied with dose condition was the one responsible for controlling random exploration which flupenthixol invariably increased.

**Figure 4.**
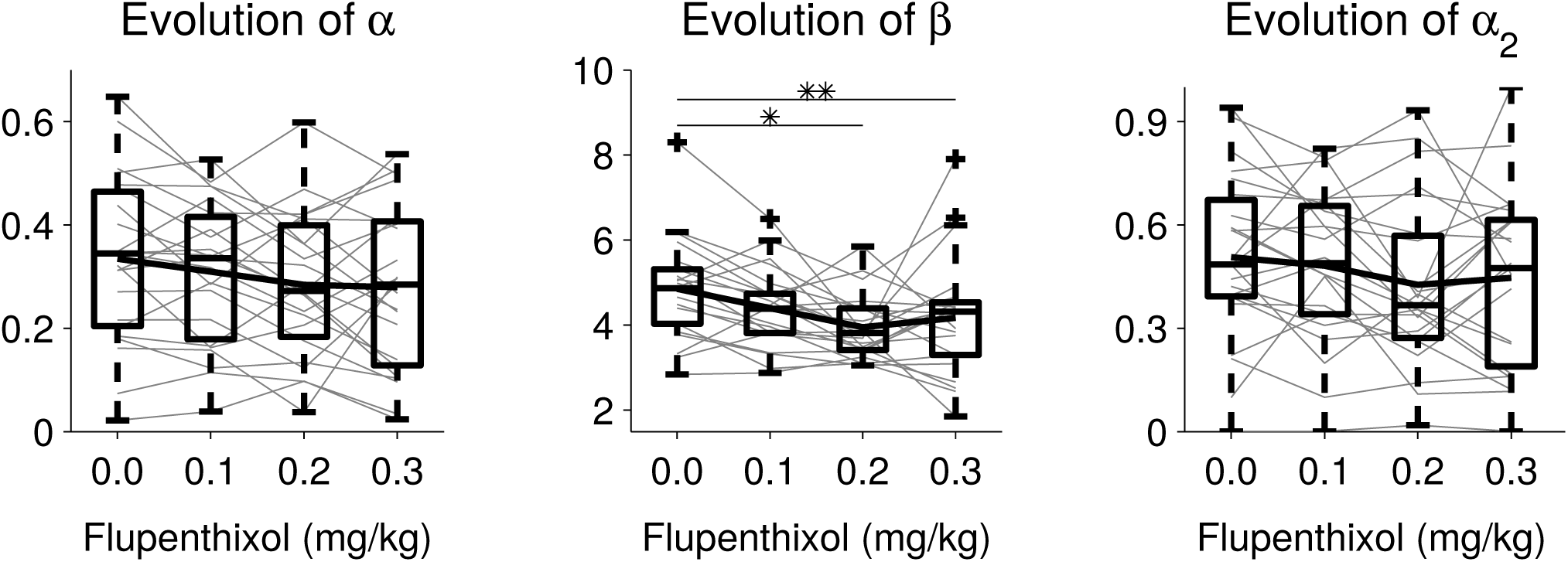
Changes of the different model parameters across flupenthixol doses in all subjects. Gray lines connect parameter values of a single individual while bold lines represent the average. Box plots represent the median, interquartile and furthest observations not considered as outliers. *: P<0.05; **:P < 0.01.

Finally, because of the strong interaction between learning rate and inverse temperature[39], we verified that our methodology was able to distinguish variations in exploration rate β from variations in the learning rate α, by applying full parameter optimization (leaving all three parameters free) to an artificial data set generated with either α or β varying between doses while the other two parameters remain constant. When only the learning rate α varied (**Fig. 5a**), the full optimization did indeed find that α significantly decreased (χ^2^(3) = 16.1, p = 0.0011), but not the two other parameters (χ^2^(3) ≤ 1.54, p ≥ 0.67). Conversely, on an artificial data set where only the exploration parameter β varied (**Fig. 5b**), the subsequent optimization correctly identified β as the only varying parameter (p = 0.0007 but p ≥ 0.12 for the other two parameters). These results clearly show that the presented computational analysis can disentangle the effects of dopamine manipulations on exploration rate from possible effects on learning rate.

**Figure 5.**
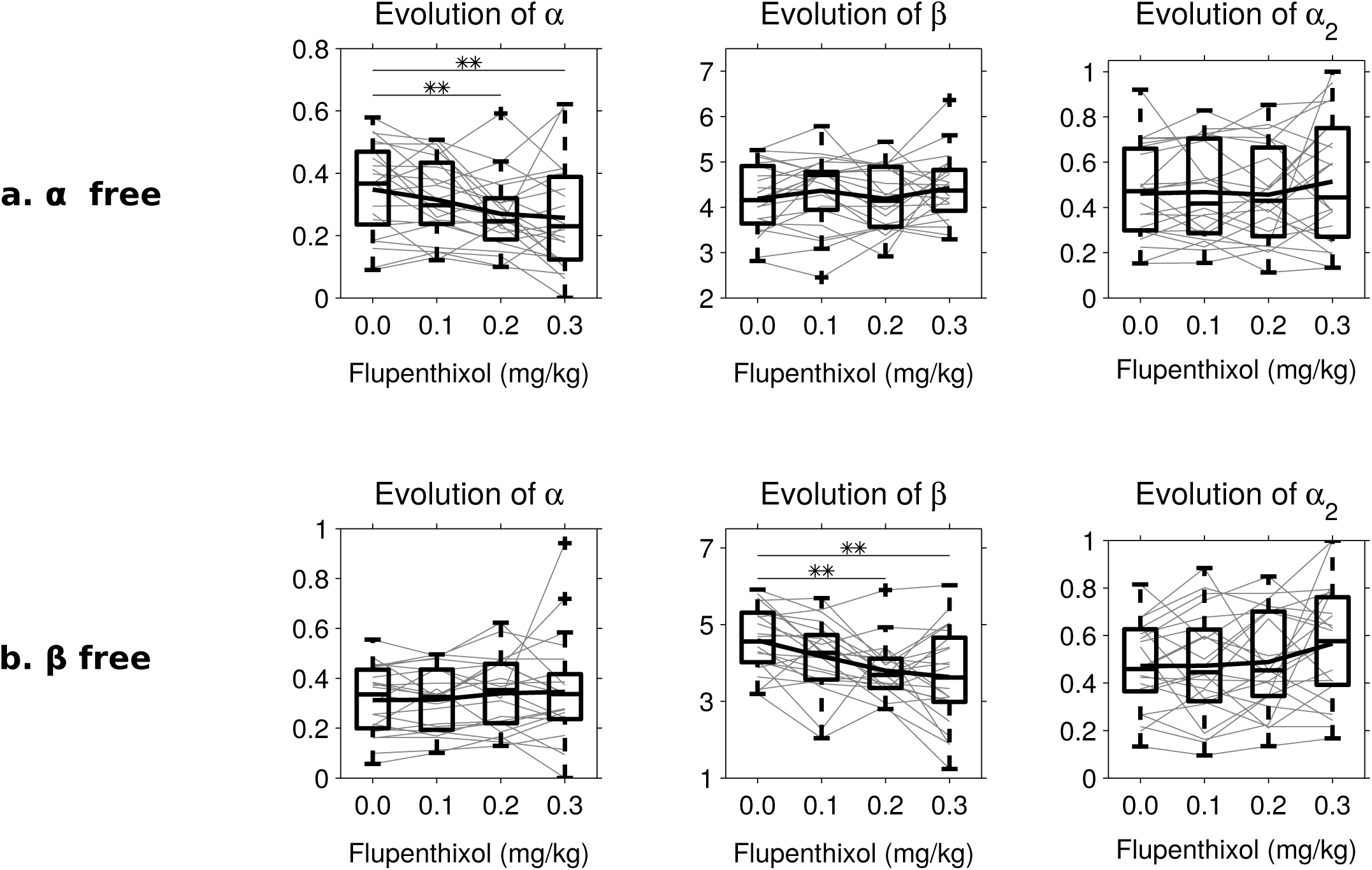
Validation of the optimization procedure. A first version of the extended Q-learning model was ed with only α allowed to vary between doses and a second version with only β allowed to vary. This ly one parameter was allowed to capture behavioural changes without compensatory adjustments her parameters. These optimized parameters were then used to generate artificial sets of data onding to a case of learning rate variation or of inverse temperature variation. We then reoptimized a model with all three parameters let free between doses as was done in the paper to ensure that only the parameter changed significantly. **(a)** Variation of the different model parameters across flupenthixol ptimized on the data set obtained from simulations where only α was free. Box plots of median, artile and furthest observations not considered as outliers which are represented as crosses. Bold resents average parameter values. An individual with an extremely high value of β was removed from the graph to make it more readable but was nonetheless included in the statistical analysis. Only α presents a significant dose effect (Friedman Anova test: χ^2^(3) = 16.1, p=0.0011) as was originally designed. **(b)** Variation of the different model parameters across flupenthixol doses optimized on the data ained from simulations where only β was free. An individual with an extreme value of β was also removed from this graph. Only β presents a significant dose effect (Friedman Anova test: χ^2^(3) = 17.1, p =0.0006).

## Discussion

This study presents a formal demonstration that in simple reinforcement learning models, a reduction of the amplitude of positive reward prediction errors directly translates into changes in random exploration levels. In rats tested on the probabilistic choice task, we experimentally show that systemic administrations of flupenthixol, a D1-D2 antagonist, dose-dependently increases random exploration, and only indirectly affects performance. Dopamine blockade increased win-shift behavior under both low and high risk conditions and noticeably late in a block when the rats had acquired the correct response. We reproduce behavioral data using unconstrained simulations and show under a variety of models that exploration rate is the only parameter significantly affected by dopamine blockade in this task.

To our knowledge, this is the first study that combines a within-subject approach with rigorous modelling to evaluate the role of dopamine levels in controlling the balance between exploration and exploitation without affecting the learning rate. The effects of various doses of dopamine antagonist in the same individual indicate that normal dopamine levels limit undirected exploration in this task and therefore bias performance toward exploitation without affecting the learning rate, a result which is consistent with that of Lee et al [34] using a different animal model, the macaque, and a different decision-making task. Undirected exploration may reflect several factors, and flupenthixol could for instance have reduced motivation[40], [41], response vigor[42] or attention. However, the observation that learning rates were unchanged argues in favor of a selective effect on exploratory choices.

In the task, performance improves within a block as one of the levers is gradually identified as target. Concurrently, win-shift decreases as learning progresses. At the beginning of each block, performance drops to chance levels or below, while win-shift increases. These high levels of win-shift in the absence of drug correspond to moments when the values of the various options have not been well identified and the rat’s uncertainty is high. Indeed, in the high risk condition, identifying the correct lever appears more difficult and is associated with both lower performance and higher win-shift levels. Under dopamine blockade, the dose-dependent increase in win-shift appears independent of uncertainty, since it does not interact with either trial rank within a block or risk level. This is consistent with the notion that dopamine unconditionally scales action values and controls noise in the last stage of decision making, where action values are converted to actual choices[29]. As to the effect of flupenthixol on lose-shift, the simulations (based on 100 runs) do show a slight increase corresponding to increased random exploration. This effect does not reach significance in the actual behavioral data, where it is probably more difficult to detect close to the ceiling value (probability of lose-shift in random choice: 0.67).

Several past studies[26], [33], [43] have focused on the effects on choice of inter-individual differences in dopaminergic function. Our data thus stand in contrast with those of Beeler *et al.[33], [43]* who observed that hyperdopaminergic mice allocate more time and energy to expensive options. On the basis of computational modeling, the authors interpreted their data as an increase in undirected exploration. However, data from primates and human subjects instead indicate increases in exploration with reduced dopaminergic activity, in agreement with our results. In particular, in an investigation of the behavioral disparities between human subjects due to genes controlling prefrontal and striatal dopamine function, Frank *et al.[26]* concluded that COMT alleles associated with lower dopamine levels increased directed exploration in their timing task. Eisenegger *et al.[35]* found in a between-subject design that a strong dose of sulpiride, a dopamine D2 receptor antagonist, in healthy human subjects favored exploration without affecting learning in a probabilistic task. These inter-individual studies supposed a fixed exploration level, and were inadequate to describe changes in exploration in the same subject. However, a similar result was achieved within subjects in an oculomotor decision task by Lee *et al.[34]* who demonstrated that injections of dopamine type 2 receptor antagonist in the dorsal striatum of two macaques deteriorated the animals’ performance in a manner best explained by an increase of noise in the decision-making process rather than an effect on learning.

In the three-arm bandit task used in the present study, we modeled learning of the correct action using an extended Q-learning model with forgetting, and we modeled choice behavior using a softmax mechanism. Our model was sufficient to account for behavioral performances as shown by i) the high similarity of the simulated (unconstrained) and experimental behavioral data (**Figs. 1, 2, 3**) and ii) the high correlation between the modeled value (constrained by the rats’ choices) of the different levers and actual choice probability, even during periods of low value when the target lever was not yet identified (**Supplementary Fig. 4**).

We show that a forgetting mechanism is required to adequately account for the rats’ behavior as a simple Q-learning mechanism appears unable to cope with the multiple target shifts (reversals) involved throughout the task (**Supplementary Fig. 5**). In our model, forgetting is important to reduce the value of competing actions even when these actions are not chosen any more, unlike simple Q-learning which only adjusts the value of actions actually performed. The observation that the forgetting rate is generally larger than the learning rate (**Fig. 4**) implies that the rats tend to persist on a choice even in the absence of reward[37]. This process stands in contrast with some theories of directed exploration[44] which predict that unchosen options become attractive as uncertainty about their outcome increases.

Our model does not include any mechanism to track uncertainty about action values, unlike several models of choice behavior in humans[4], [5], [26], [28], [45], [46]. Our simulations furthermore show that the gradual reduction in win-shift within a block does not reflect a dynamic adaptation of the model parameters since it is reproduced in the simulations where these parameters are kept constant. Instead, this decrease is a consequence of the interaction between value learning and the softmax mechanism. Choice is more variable when actions values are relatively similar, and it then becomes less variable as the values of the various actions get better differentiated. We did observe a significant effect of risk on performance and win-shift in the behavioral data, but because the same effect was present in the simulations, it is attributable to a slower acquisition of value in the high risk situation due to increased stochasticity.

As expected from the formal analysis, dopamine was found to specifically control the exploration parameter β (inverse temperature) which represents undirected exploration or random noise in the choice process converting values to actions[27], [31], [32], rather than directed exploration driven by uncertainty[5], [25], [26], [28]. Furthermore, this result is still valid with other models such as the standard Q-learning, the ε-greedy version of Q-learning and even a directed exploration model. Our data agree with the theoretical proposal by Humphries et al. (2012)[29] that tonic dopamine in the basal ganglia could modulate the exploration-exploitation trade-off during decision-making, On the basis of a prior, biologically inspired model of the basal ganglia[47], they showed that changing simulated tonic dopamine levels had similar effects as changes in the β parameter. Our study highlights a common misconception that equates the well-established role of dopamine in learning[48] with an effect on learning rate. To the extent that learning is based on reward prediction errors and action selection on a softmax mechanism, as is typically assumed in model-free reinforcement learning, our formal analysis indicates that the inverse temperature, the parameter controlling random exploration, is the only parameter affected by simple manipulations of the reward prediction error signal. Notably, in our task, both behavioral performance and fitted learning rate were largely unaffected by flupenthixol. In a probabilistic learning task, Pessiglione *et al.* [49] showed that administration of L-DOPA, a chemical precursor of dopamine known for enhancing dopaminergic functions, improved performance in accumulating gain compared to subjects under a dopamine antagonist. This improvement was attributed to an increase in learning from positive reward errors, but increased dopamine could also have reduced exploration. Similarly, Krugel 2009[36] reported that COMT alleles increasing dopamine levels were associated with better performance in a reward-based learning task with reversals, and they explained their results by a modulation of learning rate. In contrast, there are reports in humans that probabilistic learning is insensitive to dopamine antagonists[50], [51]. Our results call for careful modeling of the impact of dopaminergic manipulations in behavioral tasks as changes in random exploration rates could easily be mistaken for changes in learning rate.

As fluctuations in tonic dopamine levels track the average reward rate[24], it seems natural to use such a signal to regulate the exploration-exploitation trade-off[32]: high reward rates suggest that the current policy is appropriate and the subject could crystallize its behavior by exploiting more. Conversely, sudden drops in reward rate leading to tonic dopamine decreases may lead to increased exploration of the environment in search for better options. Dopamine levels could thus contribute to dynamically regulate exploratory choices in volatile environments where option values change with time. Here, we show that dopamine blockade affects undirected exploration independently from the changes in uncertainty levels within blocks. Dopaminergic regulation of exploration appears to occur at a longer time scale than that of a few trials, which would constitute a form of meta-learning[3], [32] adapting behavior to the general characteristics of the task rather than to immediate events.

## Methods

### Behavior

Male Long Evans rats (n=24) were obtained from Janvier Labs (France) at the age of 2 months and initially accustomed to the laboratory facility for two weeks before the beginning of the experiments. They were housed in pairs in standard polycarbonate cages (49 × 26 × 20 cm) with sawdust bedding. The facility was maintained at 21±1°C, with a 12-hour light/dark cycle (7 AM/7 PM) with food and water initially available *ad libitum*. Rats were tested only during the light portion of the cycle. The experiments were conducted in agreement with French (council directive 2013-118, February 1, 2013) and international (directive 2010-63, September 22, 2010, European Community) legislations and received approval # 5012064-A from the local Ethics Committee of Université de Bordeaux.

Animals were trained and tested in eight identical conditioning chambers (40 cm wide × 30 cm deep × 35 cm high, Imetronic, Pessac, France), each located inside a sound and light-attenuating wooden compartment (74 × 46 × 50 cm). Each compartment had a ventilation fan producing a background noise of 55 dB and four light-emitting diodes on the ceiling for illumination of the chamber. Each chamber had two opaque panels on the right and left sides, two clear Perspex walls on the back and front sides, and a stainless-steel grid floor (rod diameter: 0.5 cm; inter-rod distance: 1.5 cm). Three retractable levers (4 × 1 × 2 cm) could be inserted on the left wall. In the middle of the opposite wall, a magazine (6 × 4.5 × 4.5 cm) collected food pellets (45 mg, F0165, Bio_Serv, NJ, USA) from a dispenser located outside the operant chamber. The magazine was equipped with infrared cells to detect the animal’s visits. Three LED (one above each lever) were simultaneously lit as a signal for trial onset. A personal computer connected to the operant chambers via an Imetronic interface and equipped with POLY software (Imetronic, Pessac, France) controlled the equipment and recorded the data.

During the behavioral experiments, rats were maintained at 90% of their original weight by restricting their food intake to ~15 g/day. For pre-training, all rats were trained for 3 days to collect rewards during 30 min magazine training sessions. Rewards were delivered in the magazine on a random time 60 sec schedule. The conditioning cage was lit for the duration of each session. The rats then received training for 3 days under a continuous reinforcement, fixed ratio schedule FR1 (i.e. each lever press was rewarded with one pellet) until they had earned 30 pellets or 30 min had elapsed. At this stage, each lever was presented continuously for one session and the magazine was placed adjacent to the lever (side counterbalanced across rats). Thereafter, all three levers were on the left wall and the magazine on the right wall. The levers were kept retracted throughout the session except during the choice phases. On the next two sessions, levers were successively presented 30 times in a pseudo-random order (FR1-trials). One press on the presented lever produced a reward and retraction of the lever. On the next eight sessions, levers were presented 30 times but each time five presses were required to obtain the reward (FR5-trials). As a result, all rats readily pressed the levers as soon as they were presented. The rats then underwent 24 training sessions of training in the probabilistic choice task, 20 sessions of six trial blocks each and four double sessions of 12 blocks each.

Following 24 sessions of training in the task, rats received i.p. injections (1ml/kg) of D1-D2 receptor antagonist Flupenthixol (FLU) or saline, 20 min prior to each double session of test, for a total of 16 injections. Four doses of Flupenthixol (Cis-(Z)-Flupenthixol dihydrochloride, Sigma, dissolved in saline at 0, 0.1, 0.2 or 0.3 mg/ml) were selected according to a pilot experiment. All rats received each dose (0, 0.1, 0.2 or 0.3 mg/kg) in separate sessions according to a latin square design with at least two days of recovery between injections. After two days of recovery, a second series of injections was performed under similar conditions. In addition, on the day preceding each of the eight tests, a retraining session under saline was performed.

The experimental task (**Fig. 1a**) consisted in a three-armed bandit task where rats had to select one of three levers in order to receive the reward. A trial began with a 2 sec warning light, and then the three retractable levers were presented to the rat. Pressing one of the levers could immediately result in the delivery of a reward with various probabilities. Two different risk levels were imposed: In the low risk condition (LR) one lever was designated as the target lever and rewarded with probability 7/8 (87.5%) while the other levers were rewarded with probability 1/16 (6.25%). In the high risk condition (HR), the target lever was rewarded with probability 5/8 (62.5%) and the other two possibilities with probability 3/16 (18.75%), making discrimination of the target lever much harder. After a lever press, the levers were retracted and the trial (rewarded or not) was terminated. Inter-trial interval randomly varied in range 4.5-8 sec. Trials were grouped into unsignaled blocks of fixed length (24 trials each) characterized by a constant combination of target lever and risk. The target lever always changed between block. Therefore rats had to re-learn the target lever on each block. Blocks were ordered pseudo-randomly within a session with all combinations of target and risk counterbalanced and tested twice.

### Model fitting and simulations

Behavioral data were modeled by assuming that rats continuously assign a value to each lever. This value is adjusted according to a standard Q-learning algorithm, with the addition of a forgetting mechanism. On each trial t where an action at was chosen, values are learned gradually by first calculating a reward prediction error *δ*_*t*_ representing the discrepancy between the reward received r^t^ and what was expected, i.e. the previous estimate of the value of the chosen action *Q_t_(a_t_)*, and then using *δ_t_* to update the value of this action *Q_t+1_(a_t_)*:

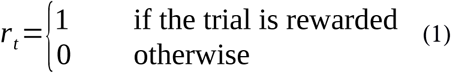

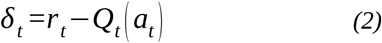

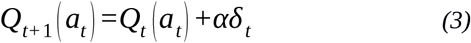

The learning rate parameter *α* determines how quickly the system learns from observed outcomes: low values ensure that action values are relatively stable. In the present task where the correct action periodically changes, higher learning rates should allow a rapid increase in performance across a block, at the cost of an increased sensitivity to the stochastic nature of reinforcement.

In order to improve the intra-block dynamics of the model, a forgetting mechanism[52] was added:

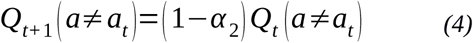

Thus, the Q-values of non-selected actions gradually regress to 0, at which they are initialized, at a rate fixed by constant *α*_2_. These values would otherwise only be updated when the corresponding action is selected. This mechanism was found to be necessary to achieve a good fit of the dynamics of win-shift (see **Supplementary Figs. 5 and 6** comparing the win-shift curves of forgetting and non-forgetting models to the experimental data) and corresponds to a perseverance mechanism independent of reward history[37]. Additionally, once combined with the action selection model described further down, we compared the log-likelihoods of the two optimized models adjusted for the number of extra parameters using either the Akaike Information Criterion (AIC) or the Bayesian Information Criterion (BIC) calculated for the whole population (n=23) and the four conditions (the better model being the one with lower scores). The Q-learning model with forgetting parameters had lower scores than the model without forgetting (AIC: 74534 *vs.* 82772; BIC: 76149 *vs.* 84386). We also compared individual log-likelihood scores for each rat and dose using the likelihood ratio test for nested models[39]. Given a model M1 nested into a more complex model M2 and their log-likelihood after optimization ll1 and ll2, d = 2*(ll2-ll1) follows a chi-squared distribution with 1 degree of freedom (1 added parameter in M2) under the null hypothesis that the log-likelihood of M2 is not better than M1. In all rats except one, the forgetting model brought a highly significant improvement in likelihood when compared to the simpler model.

Given the estimated Q-value for each action, actions are selected by sampling from a softmax probability distribution:

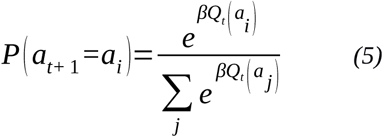

The key parameter of this function, the inverse temperature β (here called “exploration parameter”), determines the relationship between action values and action selection, or in other terms between learning and performance. Low values of β result in almost equiprobable action selection (hence exploration), independently of the learned Q-values, while high values of β greatly favour the best action over all the others (i.e. exploitation). This equation (4) is especially crucial for optimization because it defines the probability of a rat’s action at each trial given its parameter-set Θ (including learned values), which was used to calculate the likelihood of each rat’s entire history of choices, H, under the parameters of each model:

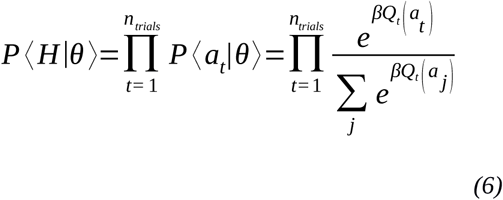

Parameter optimization was then carried out separately for each one of the four pharmacological conditions (0, 0.1, 0.2 and 0.3 mg/kg administrations of flupenthixol) by maximizing the log-likelihood of the model for each rat so as to get individual parameter-sets for each subject. Each parameter was initialized in three different points of its parameter space to avoid being trapped in local maxima.

To verify that conclusions drawn from the characteristics of the optimized model, we checked that the model was able to properly reproduce the experimental data by running unconstrained simulations using the individual parameters fitted to each rat 100 times on the full experiment. These simulations were then averaged and compared to the original data. This verification procedure has recently been advocated[53] as a standard and crucial requirement when modeling experimental data.

We used the same optimization method on four additional models. The first of these additional models is the standard Q-learning model which is identical to the model just presented except it has no forgetting mechanism. The second additional model is the forgetting Q-learning model presented earlier with an ε-greedy action selection mechanism instead of the softmax rule. This mechanism simply selects the action with highest Q-value with probability 1-ε and the remaining two actions with probability ε/2. Thus the larger the ε parameter, the more exploratory the behavior of the animal. The third model we tested is taken from the literature[25] and consists in combining Q-values, which estimate the expected payoff of a given action, with an uncertainty bonus υ, which estimates the variance of these payoffs. Q-values are updated in the same way as the extended Q-learning model (equations (2) and (3)). Each time an action a is selected and rewarded, we calculate the uncertainty prediction error ξ_t_(a) based on squared reward prediction errors (taken from equation 1) and the previous estimate of the variance:

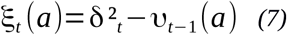

Using squared prediction errors means that it is the magnitude of reward prediction errors which matters and provides us with an estimate of variance. The expected uncertainty of the same action can then be updated using its own learning rate parameter α^φ^:

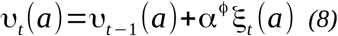

The expected uncertainties of the other two actions remain the same, and all expected uncertainties were initialized at 0, as were the Q-values. Finally, the expected uncertainties are combined with the Q-values through a weighting parameter φ before being plugged into the softmax equation:

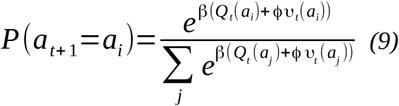

We chose this model as a representative of directed-exploration models as it can bias action selection towards potentially unrewarding choices if these are highly uncertain. As explained in the main text of this article, even when using this sophisticated version of exploration, it is still β, the parameter controlling random exploration, which is affected by dopamine inhibition.

Finally, given that our reported findings suggest the possibility of a meta-learning process based on dopaminergic control of the exploration rate, we also tested a forgetting Q-learning model in which β is controlled by an accumulation of past reward prediction errors Rt, intended to represent tonic dopamine (under the simple assumption that tonic dopamine is simply the result of past phasic activity):

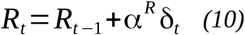

The inverse temperature is simply defined as a linear function of R_t_:

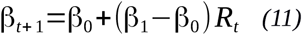

This model presents three additional parameters compared to the simple forgetting model, α^R^, which simply determines how much impact recent prediction errors have relative to older ones, β which is the value of β when Rt equals 0, and β1, the value of when Rt equals 1. Initially, we set R0 to 0 and by the same logic β equal to β0. When optimized, we found a small significant effect of flupenthixol on β1, comforting once again our results.

### Data analysis

Trials from each 24-trial block were grouped into six bins of four trials and averaged for each of the four flupenthixol doses and each of the two risk levels in each rat. Asymptotic performance and win-shift levels were estimated on the last six trials of blocks, respectively. Performance was represented by the proportion of trials where the target lever was selected. Exploration was indexed by win-shift, the proportion of trials where rats changed lever choice after being rewarded on the target lever. One rat did not complete the task under 0.3 mg/kg of flupenthixol and was therefore removed from analysis. Five other rats missed occasional parts of the blocks (no win), so statistical analysis of win-shift concerned 18 rats. Experimental or simulated data were submitted to repeated-measures ANOVAs with three factors (flupenthixol dose, risk and trial) with an additional factor (experiment/model) when appropriate. *Post-hoc* comparisons were performed using simple t-tests with Bonferroni correction. Model parameters were analyzed as a function of dose using non-parametric Friedman’s ANOVA as the distribution of β values violated the assumptions of normality (Shapiro-Wilk test of normality p<0.0001 for 0.2 mg/kg of flupenthixol), and sphericity (Mauchly test: χ^2^(5) = 24.68, p = 0.0002) required for a repeated-measures ANOVA. First-order error risk was set at 0.05 (two-sided tests).

## Acknowledgements

We thank Paul Apicella, Mark D. Humphries, Stefano Palminteri and Emmanuel Procyk for valuable discussions, and Yoan Salafranque for animal care. This work was supported by the French Agence Nationale de la Recherche (ANR) “Learning Under Uncertainty” Project under reference ANR-11-BSV4-006, “Neurobehavioral assessment of a computational model of reward learning” CRCNS 2015 Project under reference ANR-15-NEUC-0001, and by Labex SMART (ANR-11-LABX-65) Online Budgeted Learning Project supported by French state funds managed by the ANR within the Investissements d’Avenir programme under reference ANR-11-IDEX-0004-02.

## Author Contributions

A.M., M.K., E.C. and B.G. designed the experiments. V.F. carried out the behavioral protocols. V.F. and A.M. analyzed the behavioral data. M.K., B.G. and F.C. designed the models. F.C. and N.A. ran the models and simulations. F.C., M.K., B.G. and A.M. analyzed the simulation data. F.C., A.M. and M.K. wrote the manuscript. All authors discussed the results and commented on the manuscript.

## Supporting material

### Effects of positive reward prediction errors on exploration

In this section, we show mathematically that a reduction of the amplitude of reward prediction errors[54] directly translates into changes in random exploration levels in a standard Q-learning model, with or without a forgetting mechanism. In other words, we prove that inhibition of positive reward prediction errors is mathematically equivalent to changing the value of the inverse temperature and not to a change in the learning rate.

The general formula for RPE is:

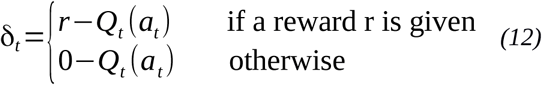

We assume that the effect of dopaminergic inhibition is to attenuate positive RPEs:

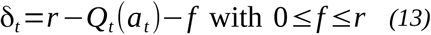

which practically results in a new reward function *r*’=*g*.*r* where 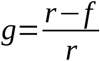 (0≤g≤1). We also assume the factor g to be constant during the learning process, which is reasonable under pharmacological or genetic manipulations if relearning is periodically required. This will change the consequences of positive RPEs (when the reward is present), but not negative RPES (in the absence of reward). Importantly, it is not equivalent to reducing learning rate, which would affect the consequences of both positive and negative RPEs. In addition, this dopaminergic manipulation is assumed not to affect the revision of value for non-selected actions if a forgetting mechanism is at play (eq. 4).

We will now show by induction that under dopaminergic inhibition, the Q-values obtained at any time during learning Q’, are just scaled-down versions of the original Q-values, Q.

Starting with Q_0_=0 and Q_0_’=0, Q_0_’=g.Q_0_ is true.

After each rewarded step of learning which only affects performed actions

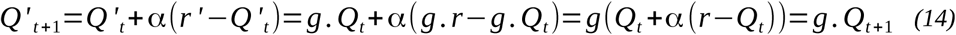

After each non-rewarded step of learning, which only affects performed actions,

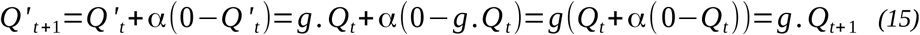

After each forgetting step, which only affects non-performed actions (if applicable)

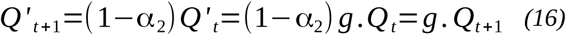

Then plugging the scaled Q-values into the softmax function we get:

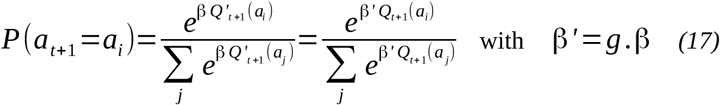

Therefore, dopaminergic inhibition, by scaling down the reward function by a factor g, is formally equivalent to multiplying the inverse temperature by the same factor g. Noticeably, although Q-values (including asymptotic values) are reduced by dopaminergic inhibition, this effect cannot be mimicked by a simple change in learning rate. Indeed, changes in learning rate do not affect asymptotic values for a constant reward function as δ_t_=0 if and only if Q_t_=r.

The same demonstration entails that increasing reward size directly reduces random exploration, thereby favoring exploitation.

**Supplementary Table 1.** Coefficients of determination of the models under various conditions. For each block, we labeled the three possible actions as either the target (action 1 in **Supplementary Fig. 4**), the previous target (action 2; for the very first block of the session, the target of the last block was used so as to ensure all lever-label combinations were equally represented) or the remaining lever (action 3). For each rat (n=23), we then averaged the experimental probabilities of each action for bins of four trials per block, and compared them to the corresponding average theoretical probabilities as determined by current Q-values plugged into the softmax function. This procedure was applied to separate doses, separate risk levels and to the entire experiment as reported in the table. In every case, the model explains a high proportion of the individual variance.

|  |  | LR | Risk<br>HR | All |
| --- | --- | --- | --- | --- |
|  | 0 | 0.9635 | 0.8412 | 0.9572 |
| Flupenthixol | 0.1 | 0.9549 | 0.8161 | 0.944 |
| Dose | 0.2 | 0.9245 | 0.8104 | 0.935 |
| (mg/kg) | 0.3 | 0.8549 | 0.677 | 0.8751 |
|  | All | 0.929 | 0.7882 | 0.9314 |

**Supplementary Figure 1.**
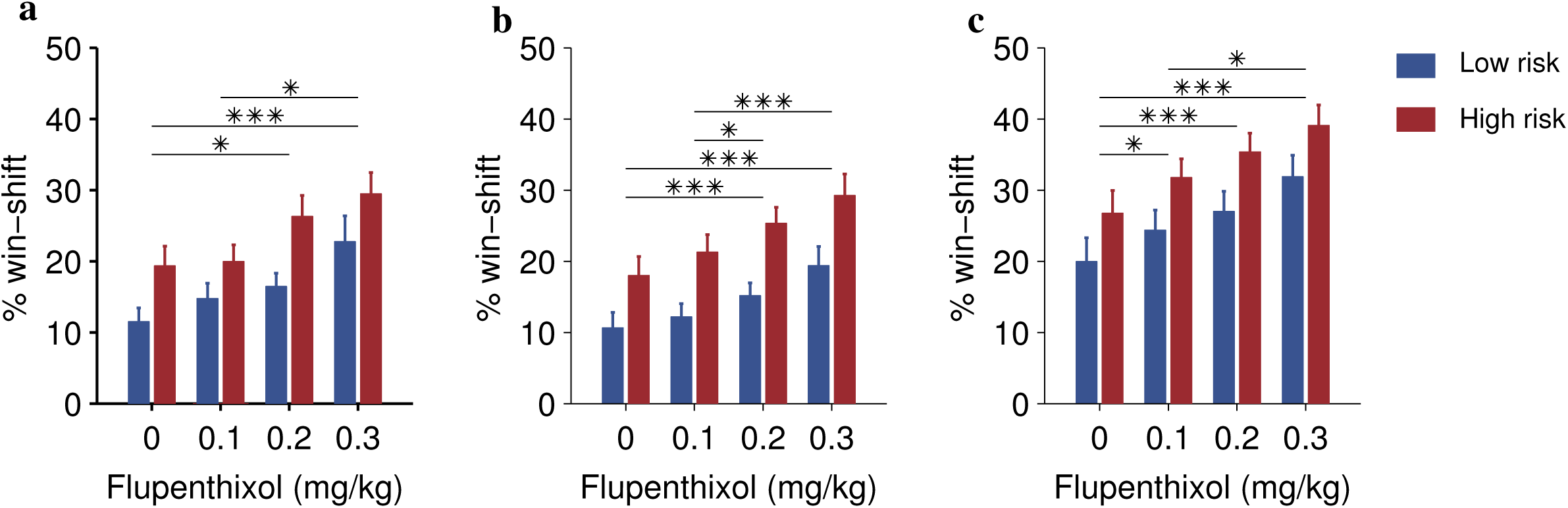
Average win-shift (mean + SEM) in the last eight trials of blocks. Post hoc tests were corrected using Bonferroni procedure (*: p < 0.05, **: p < 0.01, ***: p < 0.001). **(a)** The average win-shift
of rats (n=23) is greater in high risk blocks compared to low risk blocks (F(1,22) = 41.819, p<0.0001), and also increases with the dose of flupenthixol (F(3,66) = 13.47, p<0.0001) and by pharmacological condition (p<0.0001) suggesting that rats explore more when dopamine is inhibited. **(b)** Simulations of the extended Q-learning model replicate experimental results with similar risk (F(1,22)=165.16, p<0.0001) and flupenthixol effects (F(3,66) = 18.605, p<0.0001) on average win-shift. Furthermore, when directly confronted to experimental data using a 3-factor repeated measures Anova including a model factor, there is no significant effect of simulations (F(1,22)=1.2714, p=0.27; no significant interaction factor involving model p>0.14). **(c)** Simulations of the simple Q-learning model show significantly increased levels of win-shift compared to the original data (F(1,22) = 45.562, p<0.0001). Nonetheless, the effects of risk and dose are also captured accurately (no significant interaction between model and risk and/or dose detected, p>0.61).

**Supplementary Figure 2.**
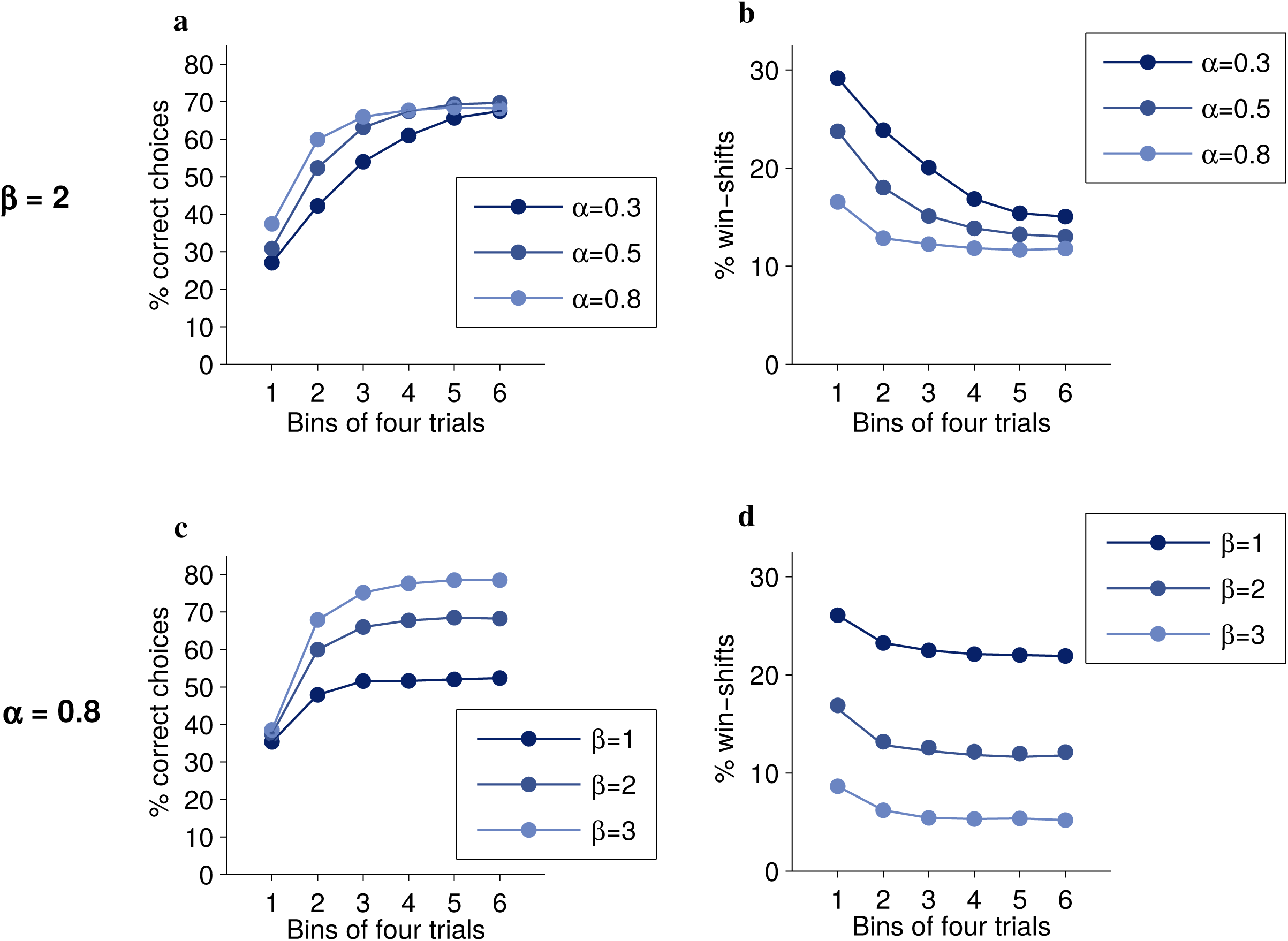
Illustration of the effects of varying either the learning rate or the exploration parameter on performance and win-shift curves based on simulations of the standard Q-learning model in low risk blocks. The range of parameters was chosen for illustrative purposes and does not reflect values actually found when the model was optimized on the experimental data. **(a)** Under different learning rates, performance curves initially increase at different rates but end up converging to similar asymptotic levels. **(b)** Win-shift curves start at different levels and decrease at different rates and eventually converge. **(c)** Under different exploration rates, performance curves converge toward different asymptotes. **(d)** Win-shift curves lie parallel to one another without ever converging.

**Supplementary Figure 3.**
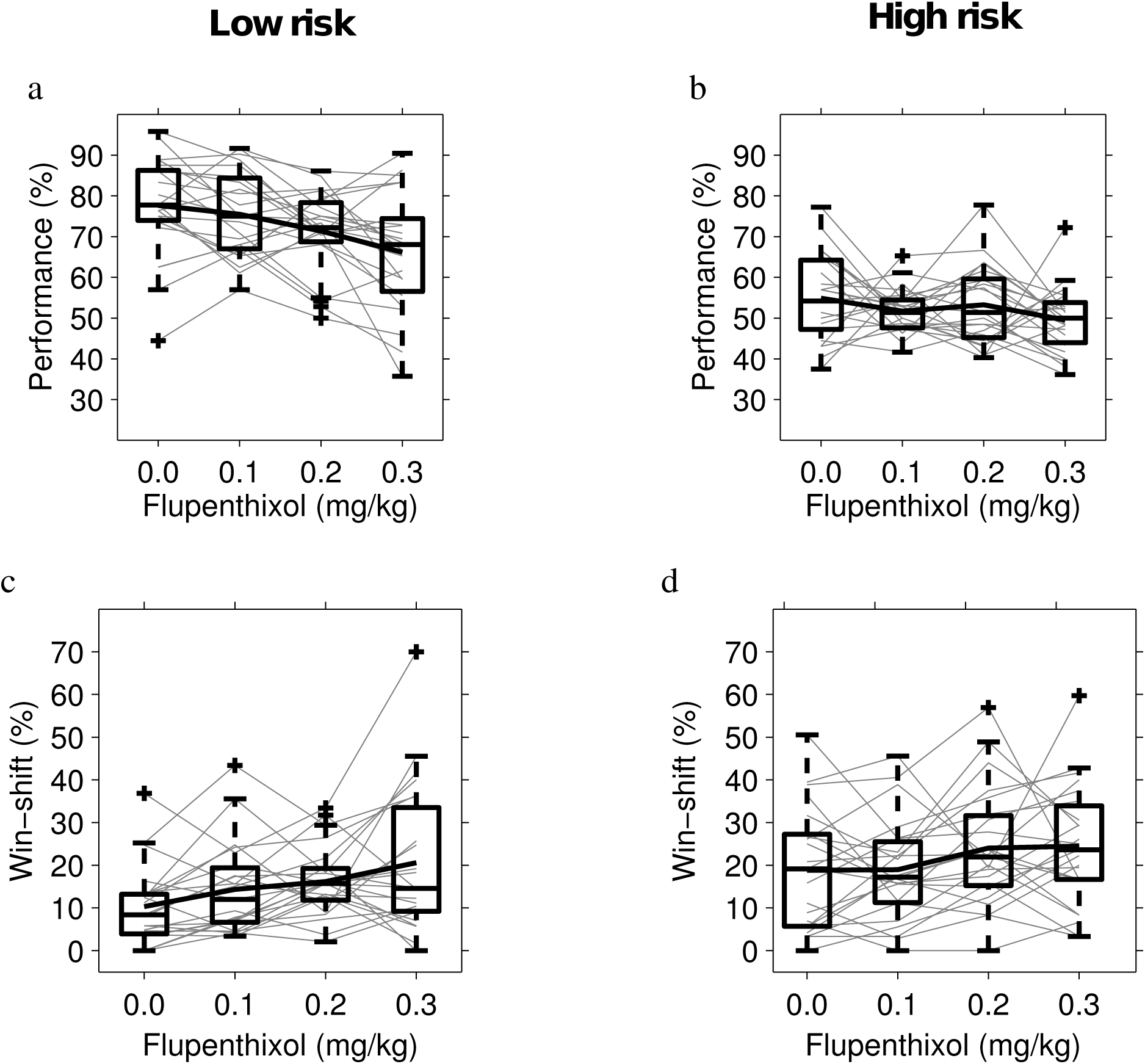
Experimental effects of flupenthixol on final performance and win-shift levels. Gray lines represent individual animals, boxes show the median, interquartile range (box edges) and most extreme data points not considered as outliers (whiskers), and crosses are suspected outliers. Bold lines represent the mean. **(a) & (b)** Effect of flupenthixol on average performance levels in the last six trials of low and high risk blocks. A significant negative effect of flupenthixol appears (F(3,66)=8.85, p<0.0001) and there is no interaction with risk (F(3,66)=1.96, p=0.13). Post hoc t-tests revealed that average performance under 0.3 mg/kg was significantly smaller than all other doses (highest p=0.020) and that average performance under 0.2 mg/kg was significantly worse than under 0 mg/kg (p=0.027). **(c)** & **(d)** Effect of flupenthixol on average win-shift in the last six trials of low and high risk blocks. A significant dose effect was detected (F(3,66)=8.76, p<0.0001) without any interaction with risk (F(3,66)=0.77, p=0.51). Post hoc t-tests showed that average win-shift increased for 0.3 mg/kg of flupenthixol when compared to 0 and 0.1 mg/kg (highest p=0.0017= and for 0.2mg/kg when compared to 0 mg/kg (p=0.0021).

**Supplementary Figure 4.**
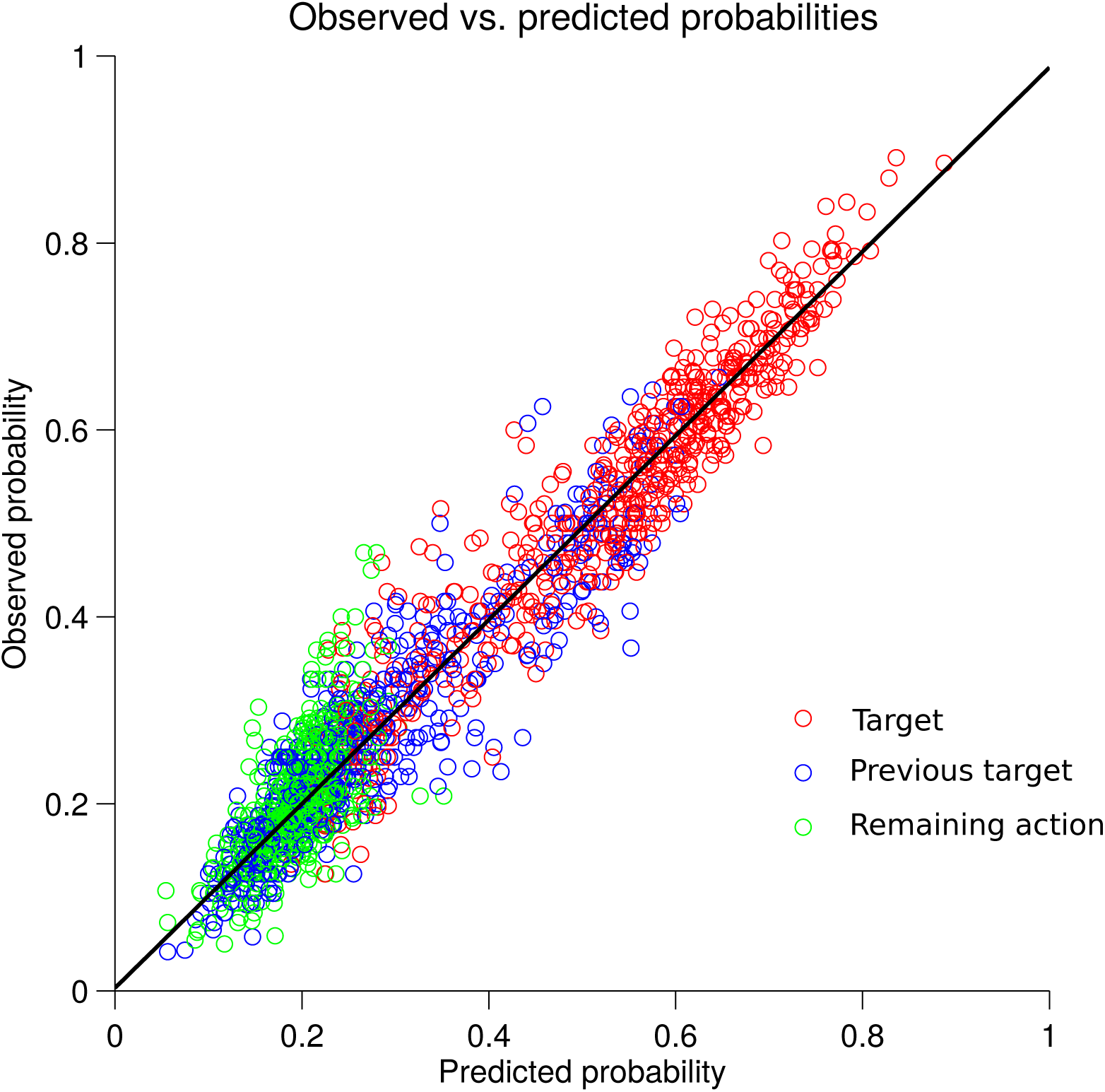
Relationship between predicted and observed probabilities of the three possible actions within a block. Predicted probability is given by the softmax function averaged in a given bin of four trials within a block. Observed probability is based on the number of times this specific action was chosen in this bin. Each data point represents the average of blocks for a given combination of rat, bin, risk level, dose and response type. Response types are identified by colors representing correct (target, red), previously correct but now incorrect (previous target, blue) or other actions (green). The same linear regression curve (y = 0.9536*x + 0.0174 is shown in black) fits all three response types.

**Supplementary Figure 5:**
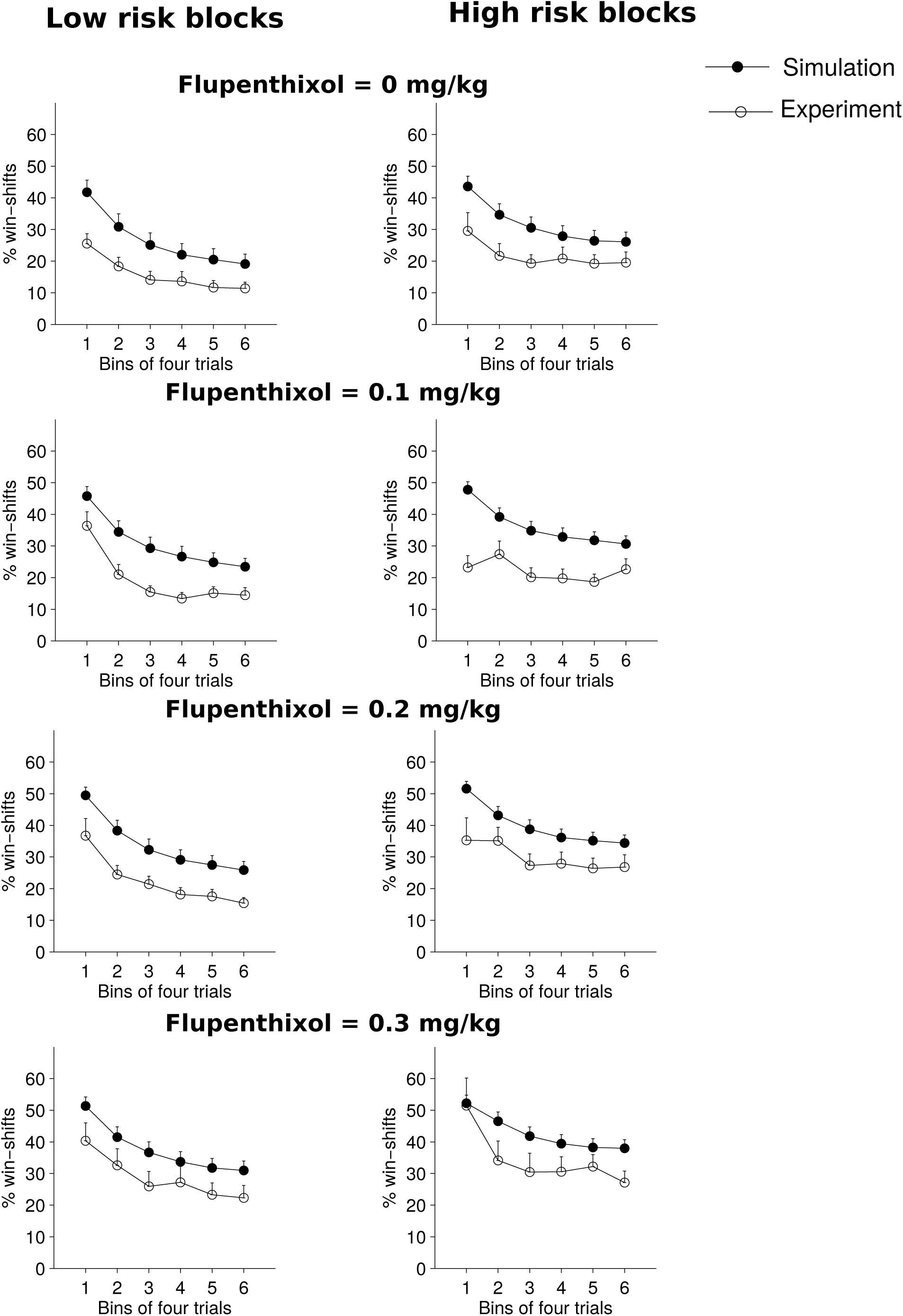
Win-shift (mean + s.e.m.) of simulations of the standard Q-learning model plotted against the experimental data for the different risk and pharmacological conditions. Contrary to the forgetting model (see **Supplementary Fig. 6**), this model was unable to reproduce this key aspect of behaviour and shifted from correct rewarded actions far more frequently than the subjects.

**Supplementary Figure 6:**
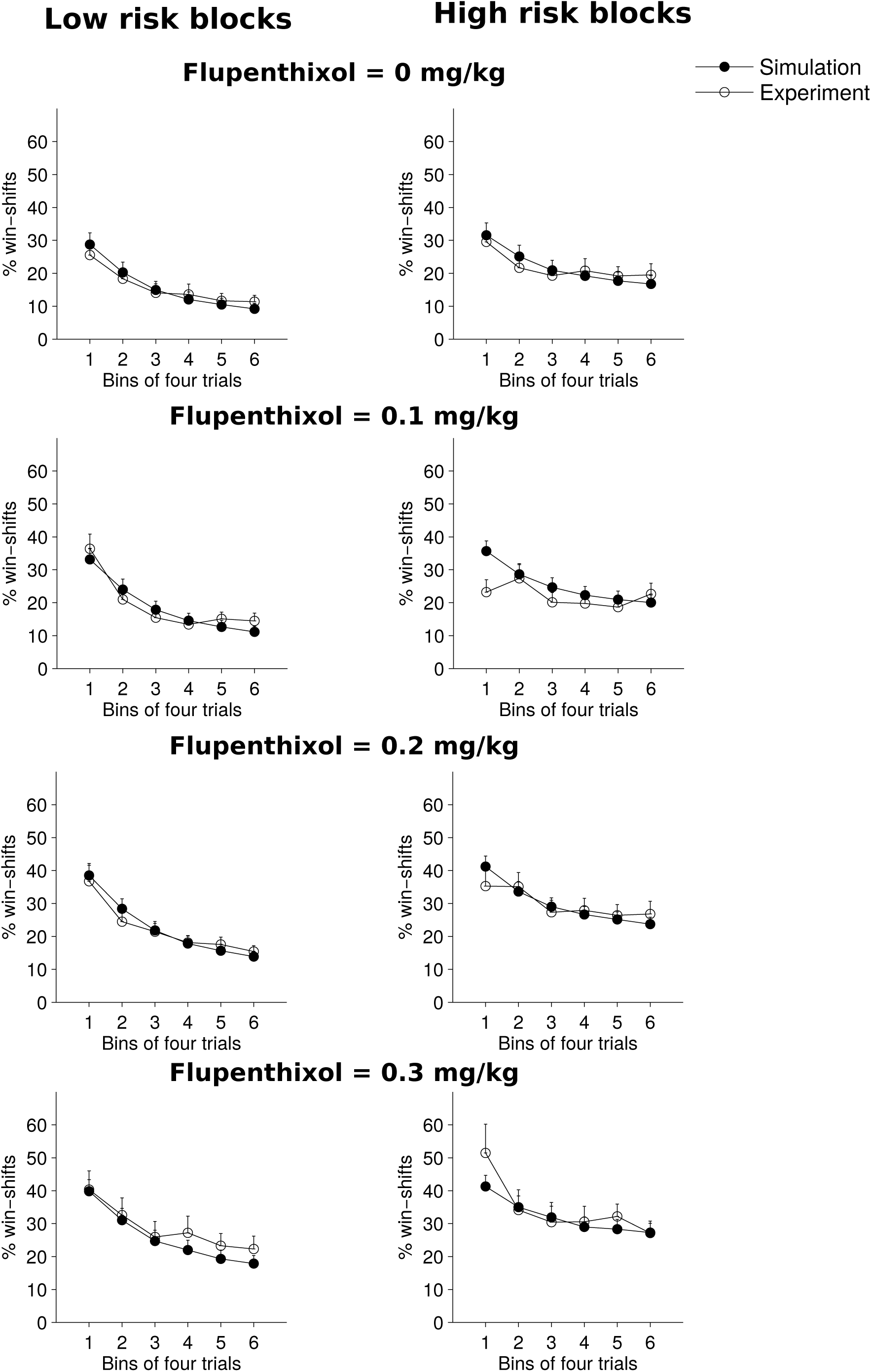
Win-shift (mean + s.e.m.) of simulations of the Q-learning model extended with a forgetting mechanism plotted against the experimental data for different doses of flupenthixol and both risk levels. This figure is in fact a different representation of **Fig. 2** for comparison with **Supplementary Fig. 5**.

**Supplementary Figure 7.**
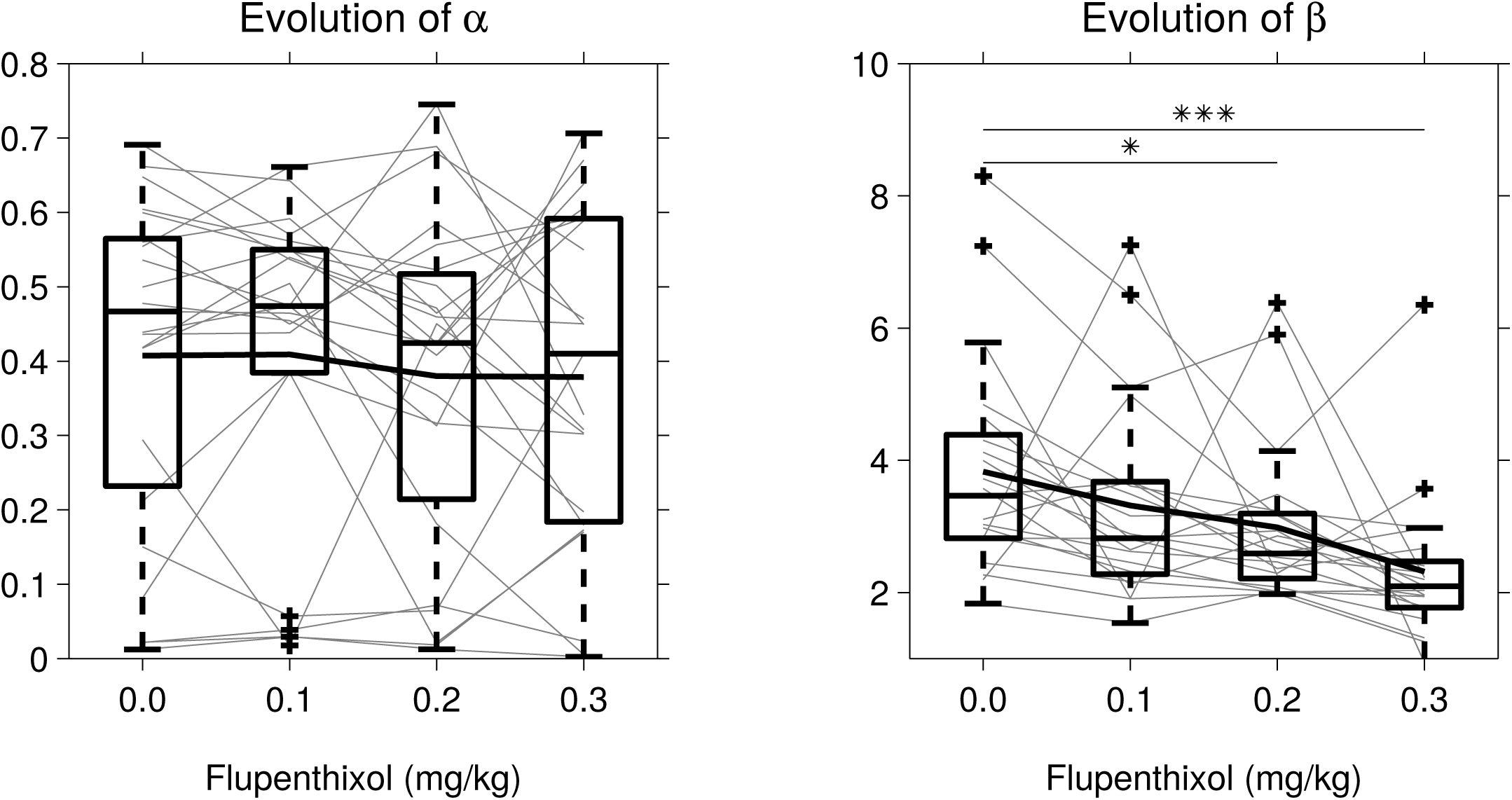
Variations of the parameters of the standard Q-learning model. Gray lines connect parameter values of a same individual. Box plots of median, interquartile and furthest values not considered as outliers represented as crosses. Bold lines plot average parameter values. As with the extended model, there is no significant dose effect on α (Friedman Anova test: χ^2^(3) = 1.90, p = 0.59) whereas β is significantly affected (χ^2^(3) = 23.5, p < 0.0001).

**Supplementary Figure 8.**
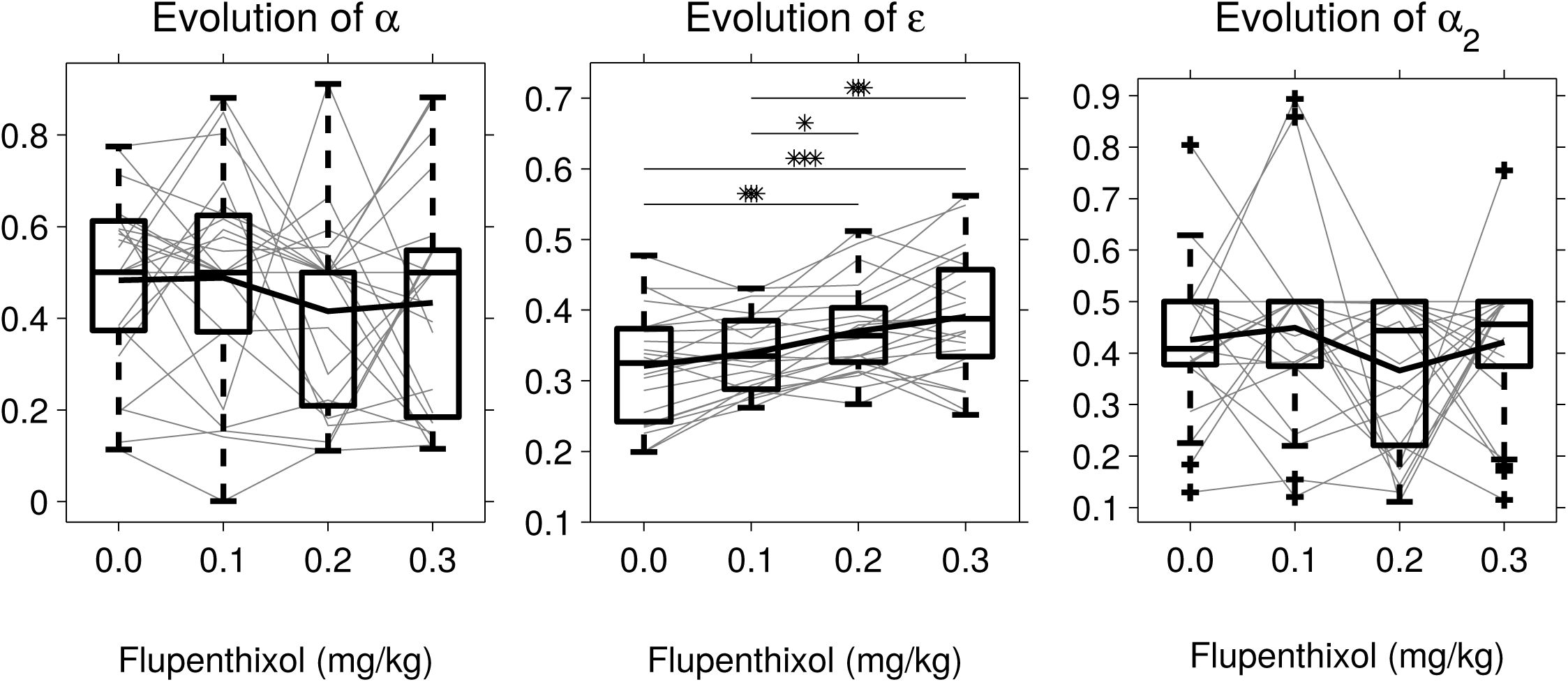
Variations of the parameters of the ε-greedy Q-learning model. Gray lines plot the parameter variations of a single individual while bold line plots average parameter values. Box plot of median, interquartile and extreme values not considered as outliers repreented as crosses. As with the softmax version of this model (**Fig. 4**), the only parameter affected by flupenthixol is the one responsible for controlling exploration, ε (Friedman Anova test: χ^2^(3)= 34.7, p<0.0001).

**Supplementary Figure 9.**
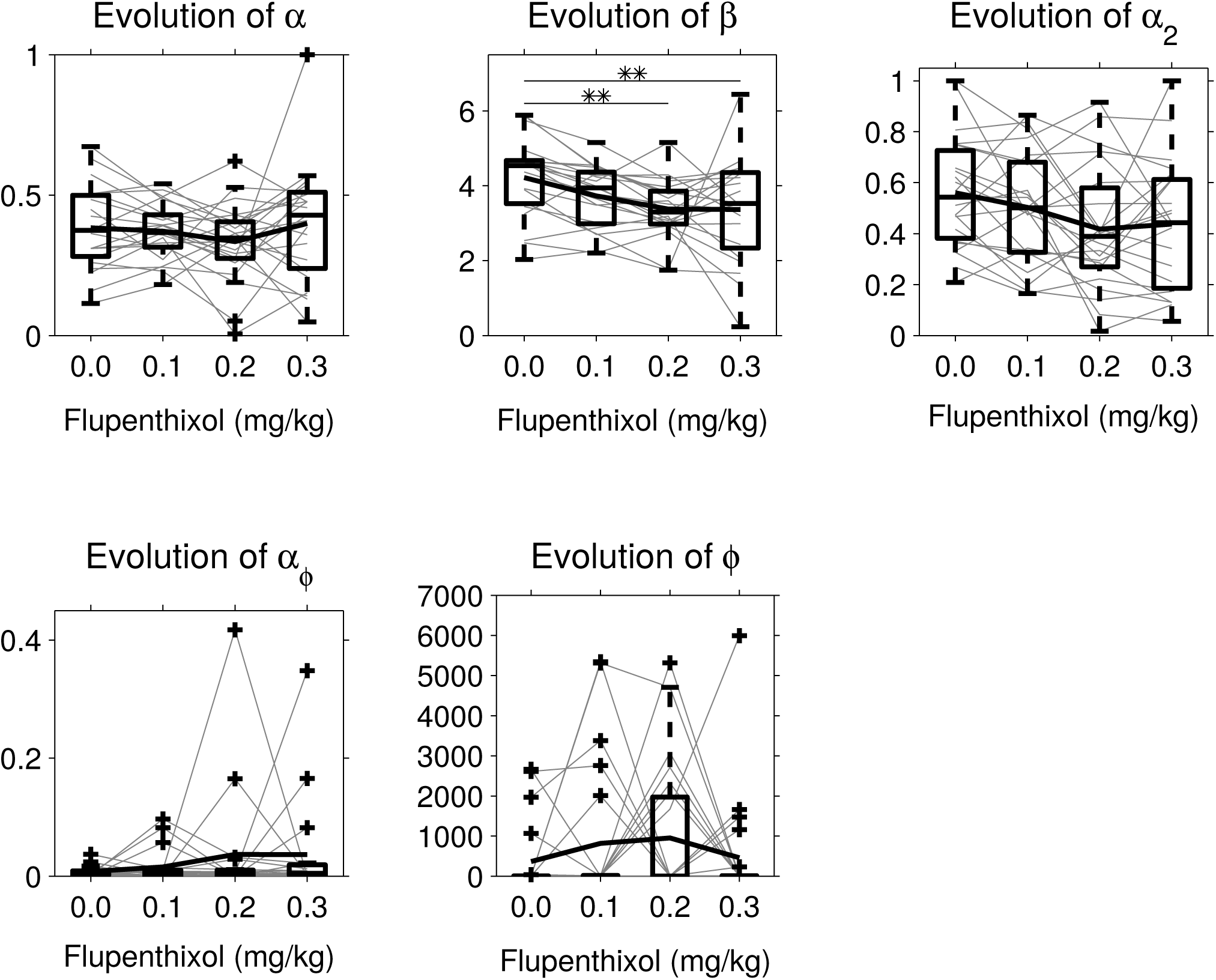
Variations of the parameters of the uncertainty bonus model. Grey lines represent single individuals and bold lines the average parameter variation. Box plots of median, interquartile and most extreme values not considered as outliers. As with previous models, the only parameter affected by flupenthixol is β (Friedman Anova test: χ^2^(3) = 16.3 p = 0.0010) which controls random exploration. Exploration targeted at uncertain options is not affected.

**Supplementary Figure 10.**
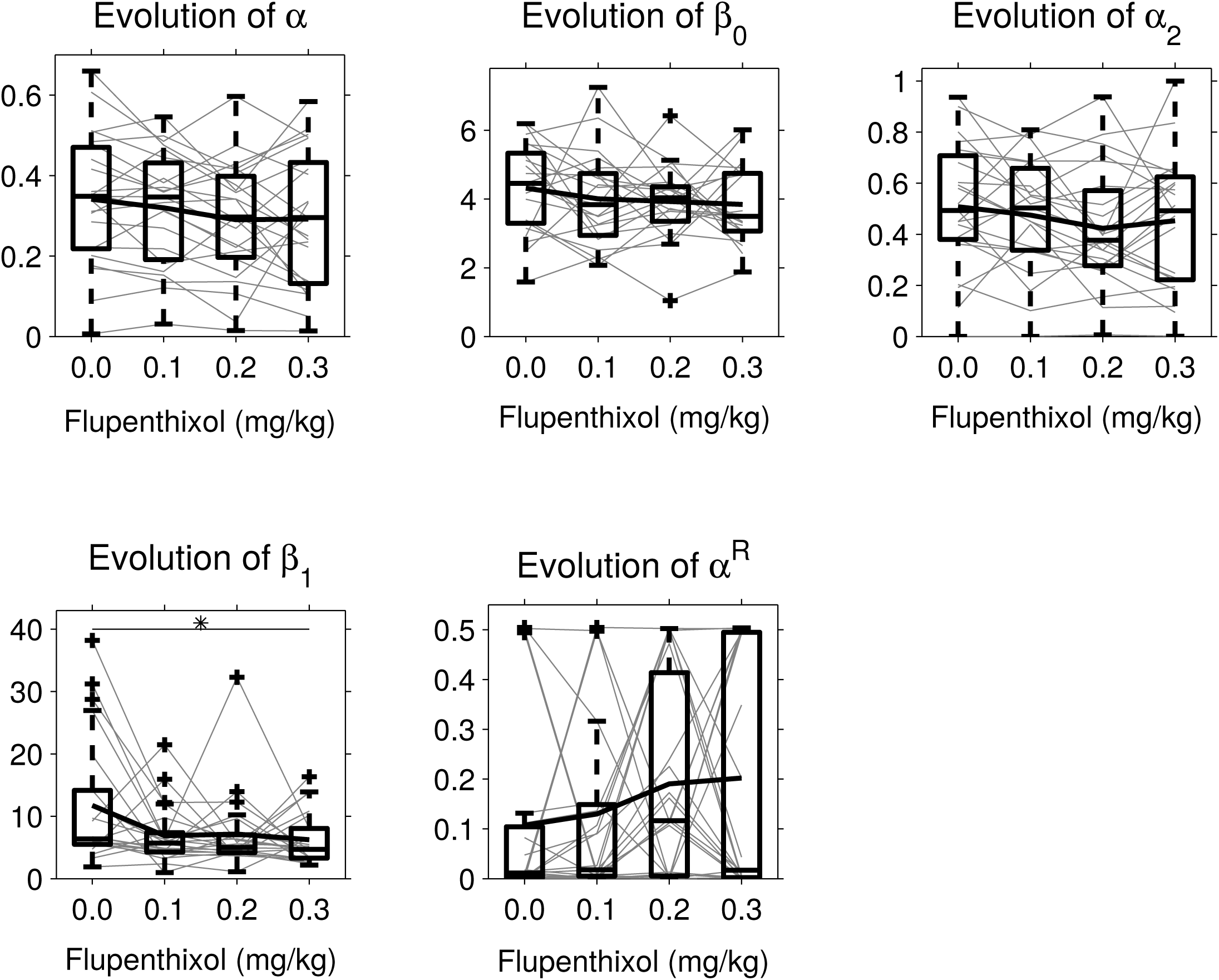
Variations of the parameters of the meta-learning model. Grey lines correspond to individual rats while the bold lines represent the average of parameter values. Box plots of median, interquartile and most extreme values which are not outliers. The only parameter affected by flupenthixol is β_1_ although the statistical effect is in fact quite modest (Friedman Anova test: χ^2^(3)= 9.3,p=0.026).

## References

[1] M. F. S. Rushworth and T. E. J. Behrens, “Choice, uncertainty and value in prefrontal and cingulate cortex,” Nature Neuroscience, vol. 11, no. 4. pp. 389–397, 2008.

[2] N. D. Daw, J. P. O’Doherty, P. Dayan, B. Seymour, and R. J. Dolan, “Cortical substrates for exploratory decisions in humans,” Nature, vol. 441, no. 7095, pp. 876–879, 2006.

[3] N. Schweighofer and K. Doya, “Meta-learning in reinforcement learning,” Neural Networks, vol. 16, no. 1, pp. 5–9, 2003.

[4] M. R. Nassar, R. C. Wilson, B. Heasly, and J. I. Gold, “An Approximately Bayesian Delta-Rule Model Explains the Dynamics of Belief Updating in a Changing Environment,” J. Neurosci., vol. 30, no. 37, pp. 12366–12378, Sep. 2010.

[5] R. C. Wilson, A. Geana, J. M. White, E. A. Ludvig, and J. D. Cohen, “Humans use directed and random exploration to solve the explore-exploit dilemma,” J exp Psychol Gen, vol. 143, no. 6, pp. 2074–2081, 2014.

[6] W. Schultz, P. Dayan, and P. R. Montague, “A neural substrate of prediction and reward.,” Science (80-.)., vol. 275, pp. 1593–1599, 1997.

[7] W. Schultz, “Updating dopamine reward signals,” Current Opinion in Neurobiology, vol. 23, no. 2. pp. 229–238, 2013.

[8] M. Watabe-Uchida, N. Eshel, and N. Uchida, “Neural Circuitry of Reward Prediction Error,” Annu. Rev. Neurosci., vol. 40, no. 1, pp. 373–394, 2017.

[9] H. M. Bayer and P. W. Glimcher, “Midbrain Dopamine Neurons Encode a Quantitative Reward Prediction Error Signal,” Neuron, vol. 47, no. 1, pp. 129–141, 2005.

[10] G. Morris, A. Nevet, D. Arkadir, E. Vaadia, and H. Bergman, “Midbrain dopamine neurons encode decisions for future action.,” Nat. Neurosci., vol. 9, no. 8, pp. 1057–1063, 2006.

[11] M. R. Roesch, D. J. Calu, and G. Schoenbaum, “Dopamine neurons encode the better option in rats deciding between differently delayed or sized rewards.,” Nat. Neurosci., vol. 10, no. 12, pp. 1615–24, 2007.

[12] M. Matsumoto and O. Hikosaka, “Two types of dopamine neuron distinctly convey positive and negative motivational signals,” Nature, vol. 459, no. 7248, pp. 837–841, Jun. 2009.

[13] D. Centonze, B. Picconi, P. Gubellini, G. Bernardi, and P. Calabresi, “Dopaminergic control of synaptic plasticity in the dorsal striatum,” Eur. J. Neurosci., vol. 13, no. 6, pp. 1071–1077, 2001.

[14] J. N. Reynolds, B. I. Hyland, and J. R. Wickens, “A cellular mechanism of reward-related learning.,” Nature, vol. 413, no. 6851, pp. 67–70, Sep. 2001.

[15] E. M. Izhikevich, “Solving the distal reward problem through linkage of STDP and dopamine signaling,” Cereb. Cortex, vol. 17, no. 10, pp. 2443–2452, 2007.

[16] V. D. Costa, V. L. Tran, J. Turchi, and B. B. Averbeck, “Dopamine modulates novelty seeking behavior during decision making,” Behav. Neurosci., vol. 128, no. 5, pp. 556–566, 2014.

[17] D. M. Haluk and S. B. Floresco, “Ventral striatal dopamine modulation of different forms of behavioral flexibility.,” Neuropsychopharmacology, vol. 34, no. 8, pp. 2041–52, Jul. 2009.

[18] S. B. Flagel et al., “A selective role for dopamine in stimulus–reward learning,” Nature, vol. 469, no. 7328, pp. 53–57, Jan. 2011.

[19] G. K. Papageorgiou, M. Baudonnat, F. Cucca, and M. E. Walton, “Mesolimbic Dopamine Encodes Prediction Errors in a State-Dependent Manner.,” Cell Rep., vol. 15, no. 2, pp. 221–8, Apr. 2016.

[20] N. L. Jenni, J. D. Larkin, and S. B. Floresco, “Prefrontal Dopamine D1 and D2 Receptors Regulate Dissociable Aspects of Decision Making via Distinct Ventral Striatal and Amygdalar Circuits.,” J. Neurosci., vol. 37, no. 26, pp. 6200–6213, Jun. 2017.

[21] J. Salamone, M. Correa, S. Mingote, and S. Weber, “Beyond the reward hypothesis: alternative functions of nucleus accumbens dopamine,” Curr. Opin. Pharmacol., vol. 5, no. 1, pp. 34–41, Feb. 2005.

[22] C. W. Berridge and A. F. T. Arnsten, “Psychostimulants and motivated behavior: Arousal and cognition,” Neurosci. Biobehav. Rev., vol. 37, no. 9, pp. 1976–1984, Nov. 2013.

[23] C. M. Stopper, M. T. L. Tse, D. R. Montes, C. R. Wiedman, and S. B. Floresco, “Overriding Phasic Dopamine Signals Redirects Action Selection during Risk/Reward Decision Making,” Neuron, vol. 84, no. 1, pp. 177–189, Oct. 2014.

[24] Y. Niv, N. D. Daw, D. Joel, and P. Dayan, “Tonic dopamine: Opportunity costs and the control of response vigor,” Psychopharmacology (Berl)., vol. 191, no. 3, pp. 507–520, 2007.

[25] J. Naudé et al., “Nicotinic receptors in the ventral tegmental area promote uncertainty-seeking,” Nat. Neurosci., no. October 2015, 2016.

[26] M. J. Frank, B. B. Doll, J. Oas-Terpstra, and F. Moreno, “The neurogenetics of exploration and exploitation: Prefrontal and striatal dopaminergic components,” in Nature Neuroscience, 2009, vol. 12, no. 8, pp. 1062–1068.

[27] W. K. Zajkowski, M. Kossut, and R. C. Wilson, “A causal role for right frontopolar cortex in directed, but not random, exploration,” Elife, vol. 6, pp. 1–18, 2017.

[28] I. Cogliati Dezza, A. J. Yu, A. Cleeremans, and W. Alexander, “Learning the value of information and reward over time when solving exploration-exploitation problems,” Sci. Rep., vol. 7, no. 1, p. 16919, Dec. 2017.

[29] M. D. Humphries, M. Khamassi, and K. Gurney, “Dopaminergic control of the exploration-exploitation trade-off via the basal ganglia,” Front. Neurosci., vol. 6, no. FEB, pp. 1–14, 2012.

[30] R. Sutton and A. Barto, “Reinforcement Learning: An Introduction,” MIT Press, Cambridge, Massachusetts, 1998. .

[31] K. Doya, “Modulators of decision making,” Nat. Neurosci., vol. 11, no. 4, pp. 410–416, Apr. 2008.

[32] M. Khamassi, P. Enel, P. F. Dominey, and E. Procyk, “Medial prefrontal cortex and the adaptive regulation of reinforcement learning parameters,” Prog Brain Res, vol. 202, pp. 441–464, 2013.

[33] J. a Beeler, N. Daw, C. R. M. Frazier, and X. Zhuang, “Tonic dopamine modulates exploitation of reward learning.,” Front. Behav. Neurosci., vol. 4, no. November, p. 170, 2010.

[34] E. Lee, M. Seo, O. Dal Monte, and B. B. Averbeck, “Injection of a Dopamine Type 2 Receptor Antagonist into the Dorsal Striatum Disrupts Choices Driven by Previous Outcomes, But Not Perceptual Inference,” J. Neurosci., vol. 35, no. 16, pp. 6298–6306, 2015.

[35] C. Eisenegger et al., “Role of dopamine D2 receptors in human reinforcement learning.,” Neuropsychopharmacology, vol. 39, no. 10, pp. 2366–75, 2014.

[36] L. K. Krugel, G. Biele, P. N. C. Mohr, S.-C. Li, and H. R. Heekeren, “Genetic variation in dopaminergic neuromodulation influences the ability to rapidly and flexibly adapt decisions.,” Proc. Natl. Acad. Sci. U. S. A., vol. 106, no. 42, pp. 17951–6, Oct. 2009.

[37] K. Katahira, “The relation between reinforcement learning parameters and the influence of reinforcement history on choice behavior,” J. Math. Psychol., vol. 66, pp. 59–69, 2015.

[38] B. B. Averbeck and V. D. Costa, “Motivational neural circuits underlying reinforcement learning,” Nat. Neurosci., vol. 20, no. 4, pp. 505–512, 2017.

[39] N. D. Daw, “Trial-by-trial data analysis using computational models,” Decis. Making, Affect. Learn. Atten. Perform. XXIII, pp. 1–26, 2011.

[40] A. Dickinson, J. Smith, and J. Mirenowicz, “Dissociation of Pavlovian and instrumental incentive learning under dopamine antagonists.,” Behav. Neurosci., vol. 114, no. 3, pp. 468–83, Jun. 2000.

[41] M. F. Barbano, M. Le Saux, and M. Cador, “Involvement of dopamine and opioids in the motivation to eat: influence of palatability, homeostatic state, and behavioral paradigms,” Psychopharmacology (Berl)., vol. 203, no. 3, pp. 475–487, Apr. 2009.

[42] Y. Niv, “Cost, benefit, tonic, phasic: What do response rates tell us about dopamine and motivation?,” Ann. N. Y. Acad. Sci., vol. 1104, pp. 357–376, 2007.

[43] J. a. Beeler, C. R. M. Frazier, and X. Zhuang, “Putting desire on a budget: dopamine and energy expenditure, reconciling reward and resources.,” Front. Integr. Neurosci., vol. 6, no. July, p. 49, 2012.

[44] B. B. Averbeck, “Theory of Choice in Bandit, Information Sampling and Foraging Tasks,” PLoS Comput. Biol., vol. 11, no. 3, pp. 1–28, 2015.

[45] T. E. J. Behrens, M. W. Woolrich, M. E. Walton, and M. F. S. Rushworth, “Learning the value of information in an uncertain world.,” Nat. Neurosci., vol. 10, no. 9, pp. 1214–21, 2007.

[46] M. Jepma, P. R. Murphy, M. R. Nassar, M. Rangel-Gomez, M. Meeter, and S. Nieuwenhuis, “Catecholaminergic Regulation of Learning Rate in a Dynamic Environment,” PLOS Comput. Biol., vol. 12, no. 10, p. e1005171, Oct. 2016.

[47] K. N. Gurney, M. Humphries, R. Wood, T. J. Prescott, and P. Redgrave, “Testing computational hypotheses of brain systems function: a case study with the basal ganglia.,” Network, vol. 15, no. 4, pp. 263–90, Nov. 2004.

[48] W. Schultz, “Neuronal Reward and Decision Signals: From Theories to Data,” Physiol. Rev., vol. 95, no. 3, pp. 853–951, 2015.

[49] M. Pessiglione, B. Seymour, G. Flandin, R. J. Dolan, and C. D. Frith, “Dopamine-dependent prediction errors underpin reward-seeking behaviour in humans,” Nature, vol. 442, no. 7106, pp. 1042–1045, 2006.

[50] T. Shiner, M. Symmonds, M. Guitart-Masip, S. M. Fleming, K. J. Friston, and R. J. Dolan, “Dopamine, salience, and response set shifting in prefrontal cortex,” Cereb. Cortex, vol. 25, no. 10, pp. 3629–3639, 2015.

[51] P. Smittenaar, H. W. Chase, E. Aarts, B. Nusselein, B. R. Bloem, and R. Cools, “Decomposing effects of dopaminergic medication in Parkinson’s disease on probabilistic action selection - learning or performance?,” Eur. J. Neurosci., vol. 35, no. 7, pp. 1144–1151, 2012.

[52] M. Ito and K. Doya, “Validation of decision-making models and analysis of decision variables in the rat basal ganglia,” J. Neurosci., vol. 29, no. 31, pp. 9861–9874, 2009.

[53] S. Palminteri, V. Wyart, and E. Koechlin, “Computational cognitive neuroscience: what is a good model of brain processes?,” pp. 1–20, 2016.

[54] F. Lesaint, O. Sigaud, S. B. Flagel, T. E. Robinson, and M. Khamassi, “Modelling Individual Differences in the Form of Pavlovian Conditioned Approach Responses: A Dual Learning Systems Approach with Factored Representations,” PLoS Comput. Biol., vol. 10, no. 2, 2014.

